# FigTreeKit: A Python toolkit for programmatic FigTree styling, taxonomy-aware clade auditing, and phylogenetic tree rendering

**DOI:** 10.64898/2026.08.27.747475

**Authors:** Zichao Zeng, Yinzhao Wang

## Abstract

FigTree is a long-standing phylogenetic tree viewer, but its GUI-centered workflow does not itself provide a versioned, batch-replayable record of styling operations. We present FigTreeKit, a Python package that serializes a supported subset of FigTree 1.4.4 annotations (!hilight, !color, and !font), audits taxonomy mappings before topology-gated clade collapse, retains selected BEAST-style metadata in the tested fixtures, and invokes a patched FigTree renderer for headless PNG, PDF, and SVG output. Across 60 independently generated balanced trees with 50–10,000 taxa (10 trees per size, each timed 10 times as technical replicates), the tree-level log–log slope of export time was 0.96 (95% confidence interval [CI], 0.91–1.01), which is compatible with approximately linear scaling over the tested range but does not prove it. The 189,801-taxon GTDB R232 bacterial reference tree was parsed and exported as a large-data scalability demonstration. On the 10,122-taxon GTDB R232 archaeal reference tree, the scripted workflow assessed 179 order-level groups; 142 multi-tip groups produced non-trivial collapses, whereas 37 singleton groups did not alter the display. The software is accompanied by 796 passing tests, a golden conformance corpus that includes acceptance tests against the bundled FigTree JAR, deterministic scenario-based topology checks, and an overall statement coverage of 81%, reported as a descriptive engineering metric. FigTreeKit is released under the GPL-2.0-or-later license as the figtreekit package on PyPI, with source code, documentation, and benchmark data archived on Zenodo.

## 1. Introduction

Phylogenetic trees are central to comparative genomics [1], molecular evolution [2], and pathogen surveillance [3]. As phylogenetic analyses increasingly involve thousands of taxa [4], reproducible and automated approaches to tree visualization are becoming increasingly important. FigTree [5] remains present in many phylogenetic workflows [6], supporting multiple layouts and extensive styling through begin figtree; … end; blocks and node annotations such as [&!hilight], [&!color], and [&!font] in NEXUS files [7]. However, the FigTree GUI does not itself provide a versioned, batch-replayable record of interactive styling operations: reapplying the same styling choices across revised or multiple datasets requires either repeated interaction or an external serialization workflow; and taxonomy-aware grouping with automated clade collapsing is not directly supported. We did not perform user-time comparisons; the limitations above concern the absence of built-in replay mechanisms rather than the feasibility of manual work.

Existing ecosystems overlap substantially with individual FigTreeKit components. Bio.Phylo [8], DendroPy [9], and ETE3 [10] provide tree input/output and metadata-bearing objects; phylotreelib [11] can write NEXUS files with selected FigTree-compatible color attributes; the standalone figtree-recolor script [12] modifies tip colors in FigTree-generated files; collapseGTDB [13] supports taxonomic-rank-based collapsing of GTDB trees; and in the R ecosystem ggtree [14] provides grammar-of-graphics-based tree visualization, whereas toytree [15] offers lightweight Python-based tree manipulation and plotting and TreeViewer [16] provides a modular GUI/CLI framework. To our knowledge, however, no maintained single Python package provides the exact combination evaluated here: serialization of the supported FigTree 1.4.4 annotations, mapping-completeness auditing, topology-gated collapse, and local rendering compatible with the patched FigTree workflow. This distinction concerns interoperability and integration with the FigTree ecosystem, not overall visualization quality or reproducibility in other ecosystems.

Several adjacent tools overlap with individual components of this workflow, including MonoPhy [17] for monophyly assessment, TaxOnTree [18] for taxonomy annotation, PhyloCloud [19] for online phylogenomic exploration, Taxonium [20] for interactive exploration of very large trees, and iTOL [21] for web-based visualization and annotation within its own ecosystem (Table 1; a more detailed tool-by-tool comparison is provided in Supplementary Section S8).

**Table 1.**
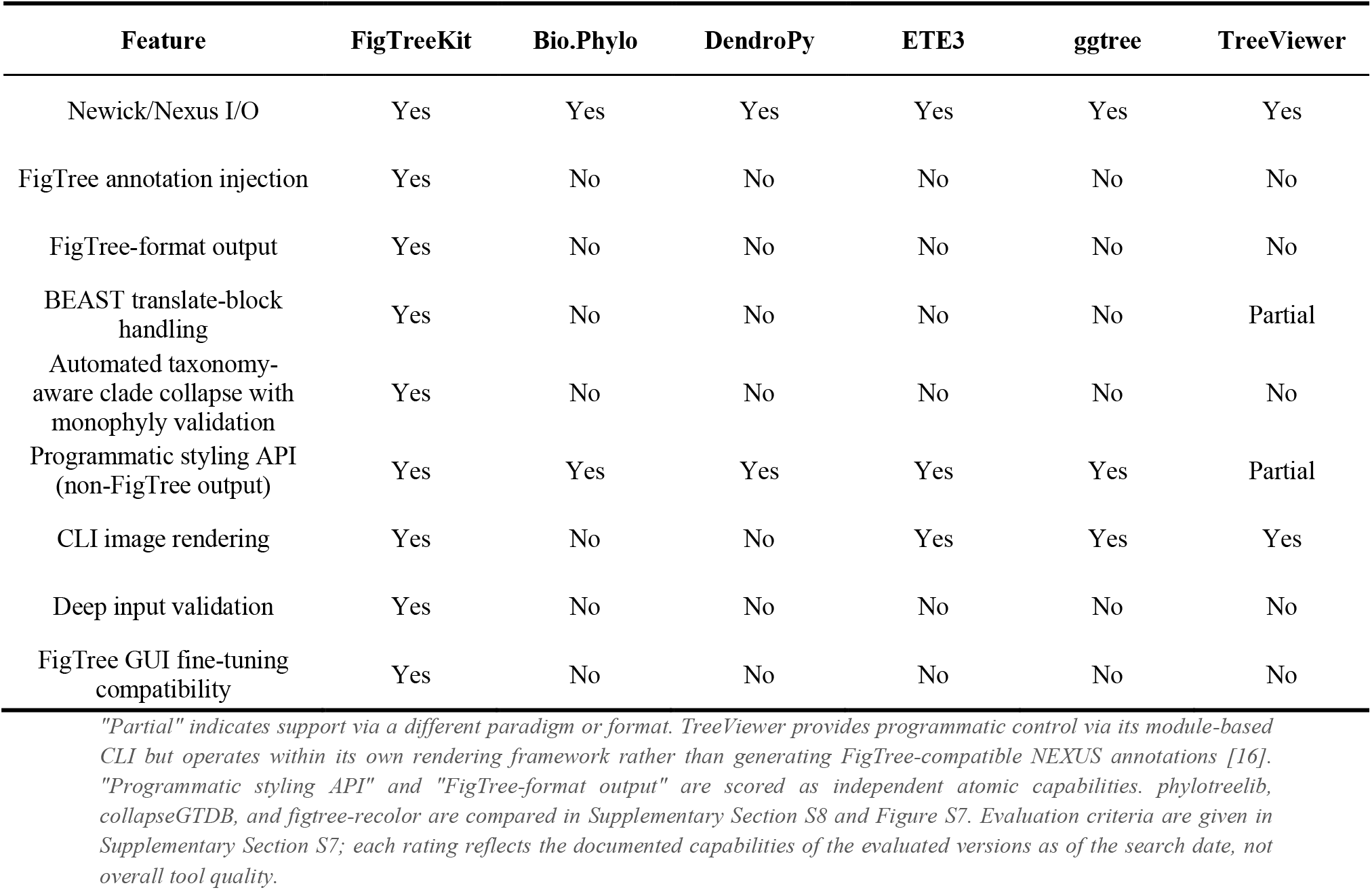
Feature comparison of phylogenetic tree visualization and manipulation tools. Features are scored according to whether the tool directly provides the specified capability within a workflow comparable to that evaluated here; the ratings are not intended to represent overall software quality. Exact FigTree 1.4.4 serialization is reported once as a specific interoperability target.

| Feature | FigTreeKit | Bio.Phylo | DendroPy | ETE3 | ggtree | TreeViewer |
| --- | --- | --- | --- | --- | --- | --- |
| Newick/Nexus I/O | Yes | Yes | Yes | Yes | Yes | Yes |
| FigTree annotation injection | Yes | No | No | No | No | No |
| FigTree-format output | Yes | No | No | No | No | No |
| BEAST translate-block handling | Yes | No | No | No | No | Partial |
| Automated taxonomy-aware clade collapse with monophyly validation | Yes | No | No | No | No | No |
| Programmatic styling API (non-FigTree output) | Yes | Yes | Yes | Yes | Yes | Partial |
| CLI image rendering | Yes | No | No | Yes | Yes | Yes |
| Deep input validation | Yes | No | No | No | No | No |
| FigTree GUI fine-tuning compatibility | Yes | No | No | No | No | No |

FigTreeKit addresses a post-inference workflow problem: it makes a tested subset of FigTree 1.4.4 styling operations scriptable, auditable, and repeatable. It does not infer phylogenies, estimate evolutionary parameters, or construct taxonomic classifications. Its principal contribution is a compatibility and workflow-integration layer for users who retain FigTree in their visualization pipeline, combining reproducible batch processing with the existing FigTree graphical interface.

FigTreeKit makes five scoped contributions, each linked to archived fixtures, expected outcomes, and test identifiers in the supplementary material: (1) conformance-tested serialization of the three supported FigTree 1.4.4 annotations (!hilight, !color, !font) over the documented input domain, validated against a golden conformance corpus; (2) parsing of the tested BEAST NEXUS TRANSLATE constructs [22], including quoted and punctuated labels; (3) deterministic reporting of conflicts between existing and injected annotations; (4) a taxonomy workflow that records mapping completeness and permits collapse only for eligible exclusive clades in the supplied rooted topology; and (5) explicit validation and error reporting for the documented malformed-input classes. FigTreeKit further integrates Bio.Phylo-based tree parsing with FigTree-compatible command-line rendering and a method-chaining API.

## 2. Materials and Methods

### 2.1 Architecture

FigTreeKit separates parsing, immutable input validation, tree-state mutation, serialization, taxonomy auditing, and optional rendering (Figure 1). The parser returns a tree plus source metadata (Newick [23] or NEXUS [7]); the FigTreeStyler records deterministic styling operations on the loaded tree state; the serializer emits FigTree-compatible NEXUS as the core output, which does not require Java; and the optional renderer consumes that serialized file to produce images through the patched FigTree JAR. The taxonomy layer supplies eligibility reports (mapping completeness and exclusive-clade verdicts) to the styler but never modifies the topology unless an explicit collapse operation succeeds. Validation and rendering failures propagate as typed exceptions (ValidationError, ExportError, RenderError).

**Figure 1.**
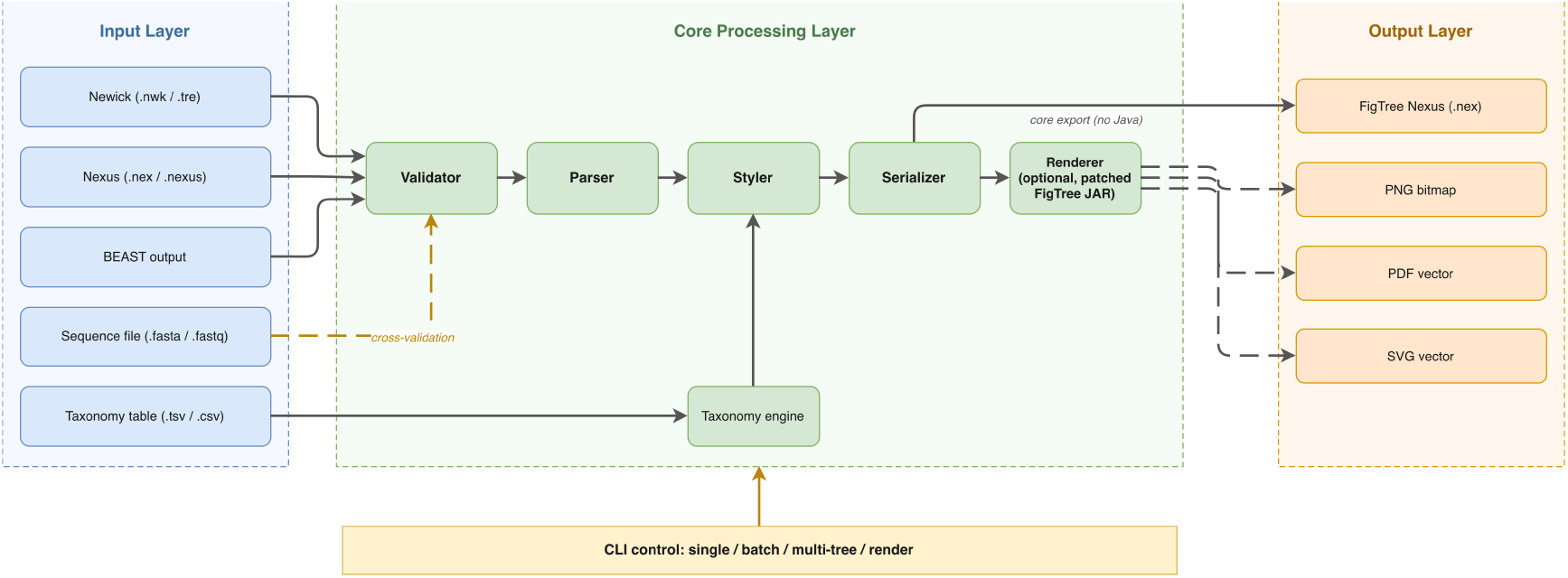
FigTreeKit architecture and data flow. Validated Newick or NEXUS input is parsed into a tree state; styling and optional taxonomy auditing annotate that state, after which the serializer emits FigTree-compatible NEXUS as the core output (no Java required). The optional patched-JAR renderer consumes the serialized file to produce PNG, PDF, or SVG output (dashed paths). Sequence files enter only through optional cross-validation; the command-line interface supports single-file, batch, multi-tree, and rendering workflows; typed failures stop the corresponding stage.

### 2.2 Core algorithms

Annotation injection. FigTreeKit emits three fully effective annotation types: !hilight (branch highlight with 3- or 5-parameter format), !color (branch color), and !font (Java Font.decode() format: Name-STYLE-size); the accepted parameter domains and units are specified in Supplementary Table S1. A fourth type, !stroke (branch width), is accepted only when the user explicitly requests it, emits a non-suppressible CompatibilityWarning on every use, and is marked ineffective because FigTree 1.4.4’s getStrokeAttribute() is an empty implementation that silently ignores the annotation; FigTreeKit never emits it implicitly. Highlight geometry and tip counts are computed from an immutable snapshot of the topology before any display collapse is applied, and collapse annotations are then registered in deterministic inner-to-outer order; this sequence preserves pre-collapse descendant counts while producing the final collapsed display (behavioral specifications with preconditions, postconditions, and test identifiers are provided in Supplementary Section S1).

Serialization rules. Values are serialized according to the observable behavior of FigTree 1.4.4’s Java createString() implementation: boolean values are written as lowercase true or false, colors as lowercase unquoted hexadecimal values, integer-valued numerical values without unnecessary decimal notation, and strings as double-quoted values. Deterministic handling of backslashes, quotation marks, structural whitespace, Unicode text, finite numeric values, duplicate FigTree keys, and conflicts between existing and injected attributes is defined in the documented grammar of Supplementary Section S1; non-finite numbers and disallowed control characters are rejected, and duplicate keys are resolved by a deterministic precedence rule reported in the operation audit. The golden conformance corpus (Supplementary Section S9) covers each of these branches.

Bracket-comment preservation. FigTreeKit parses trees without intentionally stripping bracket comments. With Biopython ≥1.80, comments are captured in Clade.comment and re-emitted during serialization, allowing the semantic content of selected BEAST-style metadata to be retained at tip, internal-node, and branch-length positions within the tested fixtures. Newly injected FigTree attributes are merged with existing comments (for example, [&support=90,!color=#00ff00]). Preservation was evaluated by comparing each selected metadata key, value, multiplicity, and clade association before and after serialization; count equality alone was not treated as evidence of preservation. Tree-level rooting directives (e.g., [&R]) were tested separately from node comments. Byte-level identity and universal preservation of arbitrary BEAST metadata are not claimed; unsupported or ambiguous cases produce a CompatibilityWarning and are listed in the machine-readable audit (Supplementary Section S9, Table S8).

State-machine parsing of translate blocks. BEAST NEXUS files may contain translate blocks that map numeric identifiers to taxon names. FigTreeKit uses an explicit four-state parser (NORMAL, IN_SINGLE_QUOTE, IN_DOUBLE_QUOTE, and ESCAPING) that treats commas as delimiters only outside quoted regions and resolves doubled-quote escapes deterministically. At end of input the parser validates the terminal state before appending the final entry; unterminated quotes, empty identifiers, and malformed trailing entries raise ValidationError. The complete function, a formal state-transition table, and executable fixtures defining doubled-quote decoding are provided in Supplementary Section S4 (Code S4.4).

Iterative node-depth calculation. FigTreeKit uses an iterative depth-first traversal to calculate root-to-node cumulative branch lengths, avoiding recursion for this operation and providing linear time complexity with respect to the number of nodes. Repeated queries are memoized per (tree, node) with a structural fingerprint guard, so cached depths are invalidated automatically after topology, rooting, or branch-length changes; failure fallbacks are never cached, and q uncached queries can require O(qn) time (Supplementary Section S1.2). Collapsed-clade triangle geometry uses a separate time-backward height calculation based on the farthest tip in the subtree; the two semantics are implemented and tested as separate functions (Supplementary Sections S1.2 and S9). MRCA search and parts of tree serialization remain dependent on recursive Bio.Phylo implementations; therefore, extremely deep trees may require an increased Python recursion limit.

### 2.3 Taxonomy-aware analysis

FigTreeKit supports two taxonomic annotation formats: embedded format A, using underscore-delimited rank prefixes (for example, _d_Bacteria_p_Cyanobacteriota_…), and tabular format B, following the GTDB/QIIME-style representation (d Bacteria;p Cyanobacteriota;…) [24]. Three delimiter-parsing modes are available: reverse (default), greedy, and segment; Supplementary Table S1 defines each grammar with worked examples for ambiguous underscores, empty ranks, duplicate identifiers, and non-standard prefixes. Custom rank prefixes can be specified globally through -- taxonomy-levels or per instance through TaxonomyMapper(prefixes=…). When embedded and tabular taxonomies disagree, the selected source-priority rule is applied explicitly and the conflict is retained in the audit report.

The taxonomy-aware workflow gates topology-gated collapse on a completeness audit. First, check_taxonomy_completeness() is executed with the same mapping and parsing configuration and identifies missing taxonomic ranks and unmapped tips; this audit precedes interpretation because each verdict is qualified as “among mapped sampled tips”. Second, analyze_taxonomy() parses all tips and assesses each taxonomic group under the implemented criterion, where “monophyletic” denotes recovery as an exclusive clade in the supplied rooted topology and does not establish biological support (e.g., bootstrap or posterior support). Third, collapse_by_group() collapses only validated exclusive clades. Non-monophyletic groups are skipped with a warning or, when --strict is enabled, cause the workflow to terminate; unmapped tips nested inside a group’s MRCA conservatively refuse the collapse. Because monophyly depends on rooting, the analysis requires a rooted tree with an appropriate outgroup, and trees without an explicitly confirmed root should not be auto-collapsed. Special identifiers such as LUCA, LACA, LBCA, and root facilitate high-rank localization. Nested collapses are performed from inner to outer regions according to region size. The complete workflow is shown in Supplementary Figure S3.

### 2.4 Validation and input hygiene

Multiple validation layers assess input integrity, including bracket balance, negative branch lengths, duplicate tip names, empty nodes, FASTA/FASTQ alphabet constraints, and tree–sequence cross-validation. Input-hygiene screening rejects control characters and Unicode bidirectional override characters (U+202A–U+202E and U+2066–U+2069); this mechanism is intended to identify anomalous input rather than to serve as a security boundary. The validation matrix specifies which findings stop loading and which permit processing with a warning: hard validation failures (ValidationError) prevent loading, whereas compatibility issues (CompatibilityWarning) allow processing to continue; negative branch lengths generate a warning in default mode and raise ValidationError only in strict mode, and unlabeled internal nodes remain valid. Rendering failures, including non-zero JVM exit codes, timeouts, and missing or invalid output files, raise RenderError, a subclass of ExportError. The validation workflow is illustrated in Supplementary Figure S4.

### 2.5 Rendering integration

FigTreeKit writes the annotated NEXUS tree to a temporary file and invokes the FigTree JAR in headless mode using the - graphic option to generate image output. Serialization compatibility and headless rendering are evaluated separately: the serialization layer targets stock FigTree 1.4.4 annotation semantics, whereas rendering uses a patched v1.4.4 JAR. The bundled patched JAR modifies four aspects of stock FigTree 1.4.4: (1) clade-collapse rendering in the radial layout (RadialTreeLayout.java); (2) a radial time axis for the polar layout (ScaleAxisPainter.java); (3) preservation of insertion order for discrete color categories (DiscreteColourDecorator.java); and (4) preservation of exact annotation colors for range fills without the stock color-brightening step (AttributableDecorator.java). The modified Java sources, build instructions, and applicable licensing information are included in the repository (_figtree_patch/and NOTICE); binary identities are pinned by SHA-256 (stock JAR 0d488f82…0346; patched JAR 13ba6b28…7892) in _figtree_patch/BUILD_PROVENANCE.md, and serialization acceptance by the stock binary is tested independently of the patched renderer.

### 2.6 Implementation

FigTreeKit was developed on macOS (Apple Silicon) using micromamba and Python 3.11; the supported deployment environments are Python 3.11 on macOS and Ubuntu, where the full test suite is executed through GitHub Actions. The core runtime dependency is Biopython (≥1.80, <2.0).

Benchmarks were performed in a separate frozen environment consisting of macOS 26.6.2, an Apple M5 processor, 32 GB RAM, Python 3.14.6, Biopython 1.87, and Java 1.8.0_501 (Supplementary benchmark_meta.json, which records the code commit); the complete test suite was additionally executed in this exact benchmark environment and passed. Windows has not been verified. The repository provides an environment.yml file pinned to Python 3.11 and a Dockerfile to facilitate deployment-environment reproduction.

## 3. Results

### 3.3 Correctness and compatibility validation

The test suite comprises 796 passing tests covering normal and error paths, regression behavior, round-trip consistency, idempotency, deep-tree validation to 1,200 levels, Hypothesis-based property tests [25], rendering output, taxonomy functions, input-hygiene scenarios, cross-validation, end-to-end CLI integration, and rendering/setup tests using controlled mocks; rendering acceptance with the bundled patched FigTree JAR is tested separately from mocked-process tests. Per-module statement coverage: _parser.py 84%, _serializer.py 91%, validators.py 86%, taxonomy.py 81%, styler.py 82%, init .py 78%, _cli.py 83%, _renderer.py 72%, _figtree_setup.py 38%; overall 81% (Supplementary Figure S1). Statement coverage is reported as a descriptive engineering metric rather than as evidence of correctness.

Beyond statement coverage, a golden conformance corpus evaluates FigTreeKit against the serialization behavior of FigTree 1.4.4. Expected annotation strings were transcribed from the FigTree 1.4.4 Java sources and pinned as independent fixtures rather than generated by FigTreeKit; the tests verify that annotation strings follow the expected FigTree serialization format, including lowercase unquoted hexadecimal colors, the supported “!hilight” parameter forms, and Java Font.decode() formatting. Round-trip tests further verify preservation of tree topology, tip sets, and branch lengths, and an acceptance test invokes the bundled patched FigTree JAR to parse the annotated output and render an image. In addition, an independent acceptance test invokes a preserved stock (unpatched) FigTree 1.4.4 binary, identified by SHA-256 in the archived release (_figtree_patch/BUILD_PROVENANCE.md), which parses the FigTreeKit-generated output and renders it without error; this binary shares no build step with the patched renderer, so acceptance is attributable to serialization compatibility rather than to the patched build. Additional tests cover clade-collapse semantics, BEAST translate blocks, and bracket-comment handling (Supplementary Section S9).

Seven deterministic scenario tests using simulated trees with known ground truth verify expected behavior for monophyletic, paraphyletic, and polyphyletic groups, as well as polytomies, incorrect rooting, and incomplete taxonomic mappings; these constitute scenario-based verification rather than an accuracy benchmark and do not estimate false-positive or false-negative rates. Validated clades are collapsed, non-monophyletic groups are rejected with reports of intruder taxa, and unmapped tips nested within a group’s MRCA conservatively prevent collapse (Table S9).

### 3.2 Performance

Across balanced random trees containing 50–10,000 taxa, export time increased approximately proportionally with tree size. Each of the 60 data points is one independently generated tree (10 trees per size; the 10 timings per tree are technical replicates summarized by the median before inference). A tree-level log–log regression of export time against taxon count yielded a slope of 0.96 (95% CI 0.91– 1.01); because the CI spans 1, the estimate is compatible with, but does not prove, linear scaling over the tested range. Peak memory increased from 0.2 to 9.0 MB across the tested tree sizes (Figure 2; Table 2). Topology sweeps using balanced, caterpillar, star, and polytomous trees showed runtime within the same order of magnitude across the tested tree shapes. Annotation-count sweeps were consistent with the predicted O(a·n) annotation-injection term reported in Table S5; when the number of annotations increased proportionally with tree size, total pipeline runtime increased super-linearly. In a serialization-overhead microbenchmark against Bio.Phylo’s basic NEXUS export, FigTreeKit required 3.6–5.0-fold more export time under the benchmark conditions (Table 3). This additional cost is associated with tree-object construction, annotation processing, and generation of FigTree-specific NEXUS blocks (Supplementary Figure S2). All measurements were generated on the frozen benchmark environment with machine-stamped provenance (benchmarks/benchmark_meta.json).

**Table 2.** FigTreeKit benchmark results on balanced random trees (10 independently generated trees per taxon count; 10 technical timing repeats per tree summarized within tree). Export time is reported as the median with interquartile range [Q1, Q3] over the 10 independent trees; peak memory is the maximum observed across the 10 trees per size (worst case).

| Taxa | Export time (s, median [IQR]) | Total time (s, median) | Peak memory (MB) |
| --- | --- | --- | --- |
| 50 | 0.0008 [0.0001] | 0.0011 | 0.2 |
| 100 | 0.0013 [0.0000] | 0.0018 | 0.3 |
| 500 | 0.0059 [0.0001] | 0.0080 | 0.6 |
| 1,000 | 0.0115 [0.0002] | 0.0154 | 1.0 |
| 5,000 | 0.0600 [0.0040] | 0.0790 | 4.9 |
| 10,000 | 0.1223 [0.0010] | 0.1605 | 9.0 |

**Table 3.** Serialization-overhead microbenchmark: export-only comparison between FigTreeKit and Bio.Phylo. This is not a feature-equivalent comparison: export times were measured after styling settings had been applied and therefore exclude annotation injection, and the benchmark is not directly comparable with the full parse–style–annotate–export workflow reported in Table 2.

| Taxa | FigTreeKit (s) | Bio.Phylo (s) | Ratio |
| --- | --- | --- | --- |
| 50 | 0.0008±0.0000 | 0.0002±0.0000 | 4.2× |
| 100 | 0.0014±0.0000 | 0.0003±0.0000 | 4.5× |
| 500 | 0.0059±0.0000 | 0.0013±0.0000 | 4.6× |
| 1,000 | 0.0120±0.0004 | 0.0025±0.0000 | 4.9× |
| 5,000 | 0.0611±0.0009 | 0.0131±0.0004 | 4.7× |
| 10,000 | 0.1258±0.0011 | 0.0263±0.0006 | 4.8× |

**Figure 2.**
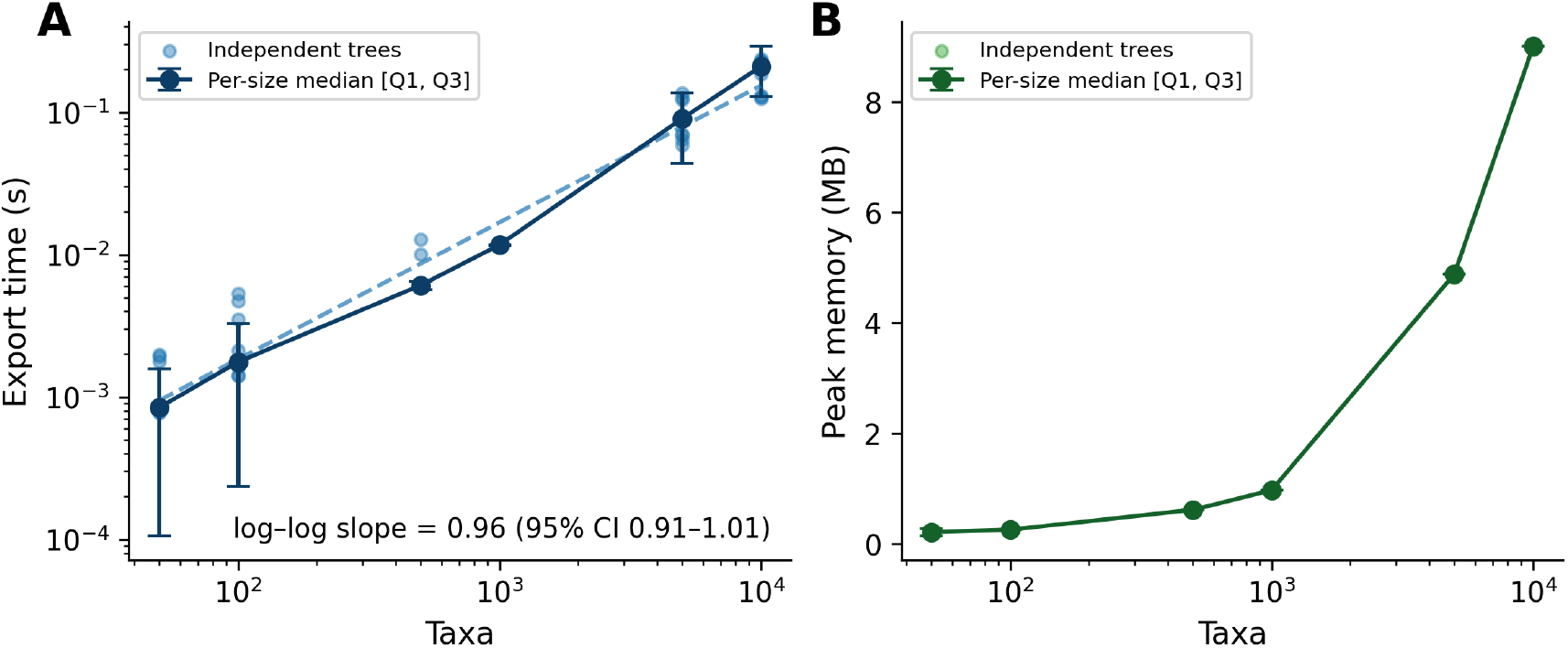
Performance of FigTreeKit on independent balanced random trees containing 50–10,000 taxa. Each light point represents one independently generated tree (median of 10 technical timing replicates); dark markers give per-size medians with interquartile ranges [Q1, Q3]. (A) Export time; the tree-level log–log regression slope is 0.96 (95% CI 0.91–1.01), compatible with approximately linear scaling over the tested range. (B) Peak memory per independent tree (process-level maximum over the parse–style–export pipeline). Benchmarks were performed in the frozen environment described in the Methods using 10 independent trees per size and 10 technical repeats per tree.

### 3.3 Large-data scalability demonstration

The GTDB R232 analyses demonstrate large-tree parsing and serialization rather than independent biological validation. The archaeal reference tree (ar53_r232.tree; 10,122 taxa) and bacterial reference tree (bac120_r232.tree; 189,801 taxa) [26–30] were successfully parsed and exported by FigTreeKit. For ar53: parse 0.06 s, export 0.16 s, peak memory 11.0 MB. For bac120: parse 1.33 s, export 4.14 s, peak memory 207.0 MB. Both trees passed validate_input_file and deep_validate_newick with no errors; because these checks are FigTreeKit’s own validators, this result demonstrates scalability and internal consistency rather than independent correctness (Supplementary Figure S6; machine-stamped raw measurements in benchmarks/gtdb_results.json, generated by the same frozen run as Tables 2–3). The bacterial tree was not interactively explored or rendered, and no large-tree rendering claim is made for that dataset; dedicated large-tree viewers such as Taxonium [20] are more suitable for interactive exploration of trees of this size (Section 4.1).

### 3.4 End-to-end workflow demonstration

To demonstrate an end-to-end workflow, we applied FigTreeKit to the GTDB R232 archaeal reference tree (10,122 taxa), including phylum-level styling and order-level clade collapsing. The completeness audit showed 100% mapping coverage (10,122 mapped tips and no unmapped tips). All 25 phylum-level groups and all 179 order-level groups were recovered as exclusive groups under the implemented criterion among mapped sampled tips. We therefore report 179 assessed order-level groups: 142 contained at least two tips and produced non-trivial visual collapses, whereas 37 singleton orders were classified as trivially exclusive (no MRCA test applies) but did not alter the display; no groups were skipped. Because the GTDB reference tree is constructed under a taxonomy-consistent framework, exclusivity at these taxonomic ranks is expected. The behavior of the workflow for non-monophyletic groups is therefore evaluated separately using the simulated ground-truth scenarios (Supplementary Table S9).

A conventional FigTree GUI workflow requires the user to locate, color, and collapse the relevant phylum-and order-level clades through repeated interaction, and produces no executable record of the styling choices unless the user maintains one externally.

The equivalent FigTreeKit workflow is implemented in a short, version-controllable script (Code 1). The workflow completes within seconds under the reported environment and provides an auditable record of the styling operations. The same script can be rerun on multiple trees with consistent styling parameters, thereby supporting reproducible batch processing without repeated manual interaction.

The demonstrated advantage is replayability within the targeted FigTree 1.4.4 pipeline: styling intent is stored in an executable script that can be version-controlled and replayed across compatible trees. We did not perform a user-time comparison, and other visualization systems provide their own reproducible workflows.

### 3.5 Rendering examples

Figure 3 illustrates the GTDB R232 archaeal reference tree styled using FigTreeKit. Panel A shows a fully expanded radial layout with phylum-level branch highlighting and coloring, whereas panel B shows a rectilinear layout with order-level clade collapse and phylum-level branch coloring. All 25 phyla use a maximum-contrast categorical palette derived from Kelly’s colors of maximum contrast [31] supplemented with Color-Universal-Design accents [32]; the shared legend applies to both panels. The inset in panel A shows a true ×2 magnification of the indicated region.

**Figure 3.**
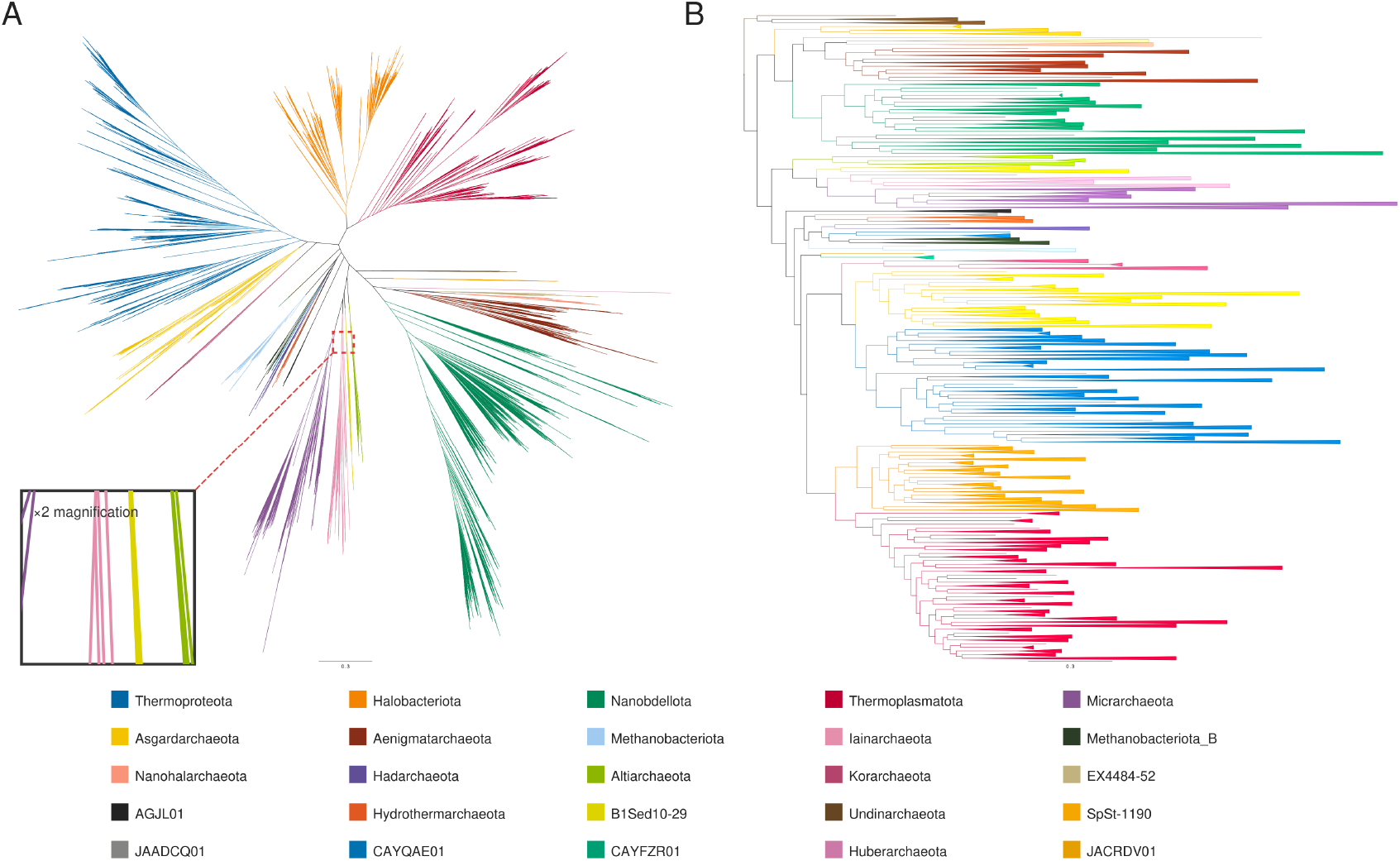
End-to-end styling of the GTDB R232 archaeal reference tree (10,122 taxa and 25 phyla) using FigTreeKit and the bundled patched FigTree JAR. (A) Fully expanded radial layout with phylum-level branch coloring and highlighting; the inset shows a 2× magnification of the indicated region. (B) Rectilinear layout after 142 non-trivial order-level collapses with phylum-level branch coloring. Branch highlighting is not applied in panel B because the tested FigTree 1.4.4 renderer exhibits a known instability when rectilinear layout, clade collapse, and highlighting are combined; the affected version, minimal reproduction, and workaround are documented in Section 4.1. The shared 25-phylum legend uses a categorical palette based on Kelly’s colors of maximum contrast [31] supplemented with Color Universal Design accents [32]; colors are separated by hue and luminance, and labels plus the legend provide non-color cues. Scale bars represent substitutions per site. Panel A is fully expanded to show tip-level detail, whereas panel B demonstrates clade-collapse behavior.

## 4. Discussion

### 4.1 Positioning and limitations

FigTreeKit is a post-inference compatibility tool, not a replacement for phylogenetic inference or for general visualization frameworks. It is most appropriate when a workflow specifically requires replayable FigTree 1.4.4 styling and later inspection in that ecosystem; the supported operating systems, annotation subset, rooting assumptions, recursive operations, and patched-renderer requirements define the present applicability boundary. FigTreeKit is not intended to replace existing phylogenetic visualization tools; rather, it targets the specific combination of FigTree ecosystem compatibility and programmatic automation. For users with established FigTree workflows, it provides a route from manual GUI-based operations to scripted, reproducible workflows while retaining compatibility with FigTree styling and BEAST-style metadata. For workflows that do not require FigTree compatibility, alternative tools such as ggtree, toytree, iTOL [21], or TreeViewer may be more appropriate. For interactive exploration of very large trees, particularly those exceeding approximately 100,000 taxa, specialized viewers such as Taxonium are better suited.

FigTreeKit is not necessarily the most suitable choice in several scenarios. First, when publication-ready figures are generated entirely within a visualization framework and no subsequent FigTree refinement is required, tools such as ggtree, iTOL, or TreeViewer may provide a more direct workflow.

Second, when many independent trees must be rendered and rendering throughput is the primary consideration, JVM startup overhead may favor tools with native or persistent rendering workflows. Third, users working exclusively within R may prefer ggtree or, for monophyly assessment alone, MonoPhy. Finally, for trees exceeding 100,000 taxa under memory-constrained conditions, full-tree loading through Bio.Phylo may become limiting.

Several limitations should be considered. First, FigTreeKit currently targets FigTree 1.4.4, whose upstream maintenance has slowed and whose Java-based rendering may become increasingly difficult to support on future operating systems. Second, bracket comments are re-emitted by the Bio.Phylo serializer in canonical positions; although their semantic content is preserved and accepted by FigTree, the resulting file may differ from the input at the byte level. Third, headless rendering depends on Java and the FigTree JAR, whereas core NEXUS export functionality does not require Java. Fourth, testing has been performed on macOS and Ubuntu using Python 3.11, with benchmarks additionally performed using Python 3.14.6; Windows and other Python versions remain unverified. Fifth, recursive MRCA operations inherited from Bio.Phylo may encounter Python recursion limits for extremely deep trees. Sixth, !stroke is accepted with an explicit compatibility warning but is ignored by FigTree 1.4.4. Finally, monophyly assessments are defined among mapped sampled tips and depend on appropriate tree rooting.

### 4.2 Long-term dependency risk on FigTree

FigTreeKit targets the specific FigTree 1.4.4 binaries evaluated in this study: serialization behavior was validated against stock FigTree 1.4.4 semantics, and rendering uses the documented patched v1.4.4 JAR. Compatibility with other FigTree releases or successor formats has not been established and will require a new conformance evaluation rather than inference from the 1.4.4 results. The Python serialization layer and the Java rendering layer are therefore versioned independently, and published workflows should record both versions. This version-specific dependency introduces several long-term risks. First, platform compatibility may become limiting because the FigTree GUI and headless renderer depend on a legacy Java runtime. This affects the optional rendering component rather than the core NEXUS-export functionality, which is implemented in Python and does not require Java. FigTree-compatible files can therefore continue to be generated independently of the rendering environment. Second, changes in future FigTree or successor formats could alter annotation semantics and require a new compatibility layer; FigTreeKit isolates serialization logic in _serializer.py, and the source modifications used to build the patched JAR are distributed in _figtree_patch/to support auditing and future maintenance. Third, if the phylogenetic visualization ecosystem moves away from FigTree, the separation between parsing, styling, and serialization provides a potential route for adapting the workflow to another backend by replacing the serialization layer.

### 4.3 Future directions

Future development will focus on several areas. The live download-and-compile workflow in _figtree_setup.py is currently experimental (statement coverage 38%) and is excluded from the supported installation route until end-to-end fixtures are available. Planned non-blocking extensions include extending continuous integration to Windows, implementing incremental serialization to reduce repeated tree parsing, developing native Python rendering to reduce the Java dependency, supporting future FigTree-compatible formats if they become available, and integrating directly with GTDB and NCBI taxonomy resources.

## CRediT author contributions statement

Zichao Zeng: Conceptualization, Methodology, Software, Validation, Formal analysis, Visualization, Writing – original draft. Yinzhao Wang: Supervision, Funding acquisition, Writing – review & editing.

## Data availability statement

FigTreeKit is free software distributed under the GPL-2.0-or-later license. The source code, test suite, conformance fixtures, benchmark scripts and raw data, figure-generation scripts, environment files, and documentation are available from the project repository (https://github.com/ZengZichao/FigTreeKit) and archived on Zenodo [33]; the archived release corresponds to the v1.1.1 tag (version-specific DOI in reference [33]; concept DOI 10.5281/zenodo.22043258). The package is distributed on PyPI as figtreekit (version 1.1.1). The example tree used in the BEAST-style workflow in Code 1 is archived with the repository under examples/data/together with a provenance README; its topology, node ages, and annotations were retained while tip labels were reformatted into GTDB embedded format A, as documented in Supplementary Table S11. The bundled rendering JAR includes the iText library; the applicable licensing information is provided in the repository NOTICE file. The GTDB R232 reference trees analyzed here are publicly available from the Genome Taxonomy Database [30] and are not redistributed. No new biological data were generated.

## Declaration of generative AI and AI-assisted technologies in the manuscript preparation process

During the preparation of this work, the authors used generative AI-assisted tools to support software development and to polish the English writing of the manuscript. All AI-assisted output was reviewed and edited by the authors; code changes were inspected, tested, and validated against the documented fixtures before inclusion. The authors take full responsibility for the content of the article and the released software.

## Code

**Code 1.**
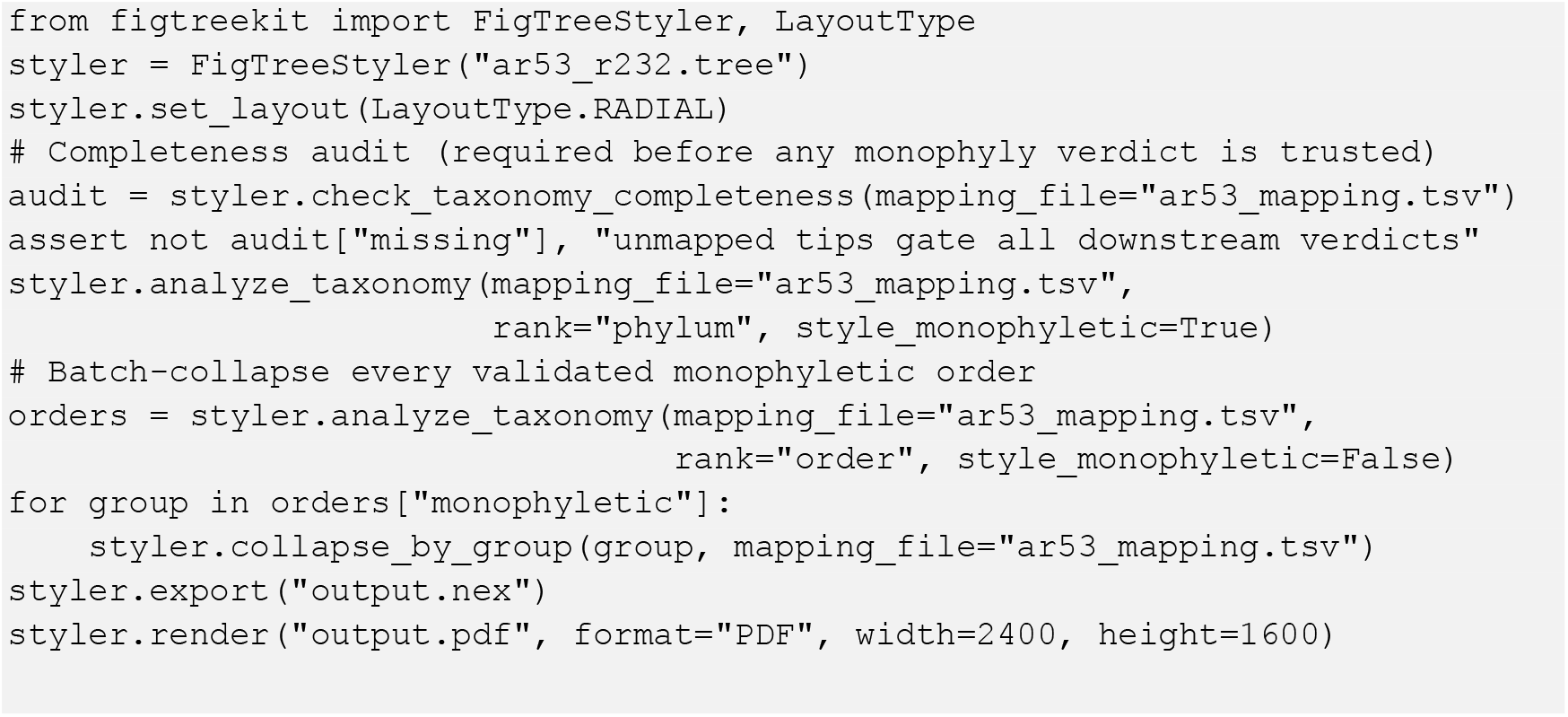
Abbreviated API illustration of the end-to-end FigTreeKit workflow for styling and collapsing the GTDB R232 archaeal reference tree. Code 1 demonstrates the audit-gated sequence (completeness audit consumed as a gate, monophyly assessment, collapse of validated order-level groups, export, rendering); it does not by itself reproduce both panels of Figure 3. The complete figure-generation workflow — including the metadata reduction step, the audit counters, and the generation of both Figure 3 panels from independently styled copies of the same input after the shared completeness audit — is examples/05_gtdb_workflow.py in the archived release.

*ar53_mapping*.*tsv is the two-column reduction (accession, gtdb_taxonomy) of the public GTDB metadata; the complete, runnable script — including the reduction step, the completeness audit, and the audit counters — is examples/05_gtdb_workflow*.*py in the project repository. A second workflow demonstrates the same pipeline on the published time-calibrated 700-tip archaeal–bacterial phylogeny of Moody et al. [34], supplied as BEAST-style annotated NEXUS: we renamed its tip labels into GTDB embedded format A to exercise the taxonomy-aware workflow, leaving the topology, node ages, and the 699 [&95%={lo, hi}] node-age 95% highest-posterior-density (HPD) interval annotations unchanged (examples/06_beast_laca_workflow*.*py). Per-clade audit values for this workflow are given in Supplementary Table S11. The GTDB tree and metadata required by the first workflow are not redistributed here; download GTDB R232 from the Genome Taxonomy Database [30] and pass the file paths as arguments (examples/05_gtdb_workflow*.*py prints download instructions when the files are absent)*.

## Acknowledgements

The authors thank the developers of FigTree (Andrew Rambaut), Biopython, and the Genome Taxonomy Database for making their software and data openly available.

## Supplementary Materials

Supplementary Section S1. Behavioral specifications for all core algorithms (preconditions, postconditions, and test identifiers).

Supplementary Section S2. Key parameter reference (Tables S1–S2).

Supplementary Section S3. Robustness test scenarios (Table S3; 28 cases).

Supplementary Section S4. Core algorithm implementation code (Code S4.1–S4.7; frozen tag).

Supplementary Section S5. FigTreeStyler class architecture (Table S4).

Supplementary Section S6. Complexity analysis (Table S5).

Supplementary Section S7. Functional comparison evaluation criteria (Table S6).

Supplementary Section S8. Tool-by-tool comparison with adjacent software (Table S7).

Supplementary Section S9. Correctness and compatibility validation (Tables S8–S11).

Supplementary Figure S1. Per-module test coverage distribution.

Supplementary Figure S2. FigTreeKit versus Bio.Phylo export performance.

Supplementary Figure S3. Taxonomy-aware analysis workflow.

Supplementary Figure S4. Input validation and hygiene-screening workflow.

Supplementary Figure S5. Pipeline-stage breakdown and topology sweep.

Supplementary Figure S6. GTDB R232 real-dataset validation.

Supplementary Figure S7. Extended tool-comparison heatmap.

## References

Koonin, E.M., Wolf, Y.I., 2008. Genomics of bacteria and archaea: the emerging dynamic view of the prokaryotic world. Nucleic Acids Res. 36, 6688–6719. 10.1093/nar/gkn668.

Yang, Z., Rannala, B., 2012. Molecular phylogenetics: principles and practice. Nat. Rev. Genet. 13, 303–314. 10.1038/nrg3186.

Bedford, T., Riley, S., Barr, I.G., Broor, S., Chadha, M., Cox, N.J., Daniels, R.S., Gunasekaran, C.P., Hurt, A.C., Kelso, A., Klimov, A., Lewis, N.S., Li, X., McCauley, J.W., Odagiri, T., Potdar, V., Rambaut, A., Shu, Y., Skepner, E., Smith, D.J., Suchard, M.A., Tashiro, M., Wang, D., Xu, X., Lemey, P., Russell, C.A., 2015. Global circulation patterns of seasonal influenza viruses vary with antigenic drift. Nature 523, 217–220. 10.1038/nature14460.

Delsuc, F., Brinkmann, H., Philippe, H., 2005. Phylogenomics and the reconstruction of the tree of life. Nat. Rev. Genet. 6, 361–375. 10.1038/nrg1603.

Rambaut, A., 2018. FigTree v1.4.4 [software]. http://tree.bio.ed.ac.uk/software/figtree/ (accessed July 2026).

Theys, K., Lemey, P., Vandamme, A.-M., Baele, G., 2019. Advances in visualization tools for phylogenomic and phylodynamic studies of viral diseases. Front. Public Health 7, 208. 10.3389/fpubh.2019.00208.

Maddison, D.R., Swofford, D.L., Maddison, W.P., 1997. NEXUS: an extensible file format for systematic information. Syst. Biol. 46, 590–621. 10.1093/sysbio/46.4.590.

Cock, P.J.A., Antao, T., Chang, J.T., Chapman, B.A., Cox, C.J., Dalke, A., Friedberg, I., Hamelryck, T., Kauff, F., Wilczynski, B., de Hoon, M.J.L., 2009. Biopython: freely available Python tools for computational molecular biology and bioinformatics. Bioinformatics 25, 1422–1423. 10.1093/bioinformatics/btp163.

Sukumaran, J., Holder, M.T., 2010. DendroPy: a Python library for phylogenetic computing. Bioinformatics 26, 1569–1571. 10.1093/bioinformatics/btq228.

Huerta-Cepas, J., Serra, F., Bork, P., 2016. ETE 3: reconstruction, analysis, and visualization of phylogenomic data. Mol. Biol. Evol. 33, 1635–1638. 10.1093/molbev/msw046.

Pedersen, A.G., 2026. phylotreelib: Python library for phylogenetic tree formats, including NEXUS output with FigTree-compatible color annotations (Version 2.3.0) [software]. PyPI. https://pypi.org/project/phylotreelib/ (accessed July 2026).

acorg, 2017. figtree-recolor: a script to re-color a FigTree saved Nexus file [software]. GitHub. https://github.com/acorg/figtree-recolor (accessed July 2026).

genomewalker, 2022. collapse-gtdb-tree: a tool to collapse a GTDB tree to a certain taxonomic rank [software]. GitHub. https://github.com/genomewalker/collapse-gtdb-tree (accessed July 2026).

Yu, G., Smith, D.K., Zhu, H., Guan, Y., Lam, T.T.-Y., 2017. ggtree: an R package for visualization and annotation of phylogenetic trees with their covariates and other associated data. Methods Ecol. Evol. 8, 28–36. 10.1111/2041-210X.12628.

Eaton, D.A.R., Matschiner, M., 2020. toytree: a minimalist tree visualization and manipulation library for Python. Methods Ecol. Evol. 11, 187–191. 10.1111/2041-210X.13313.

Bianchini, G., Sánchez-Baracaldo, P., 2024. TreeViewer: flexible, modular software to visualise and manipulate phylogenetic trees. Ecol. Evol. 14 (2), e10873. 10.1002/ece3.10873.

Schwery, O., O’Meara, B.C., 2016. MonoPhy: a simple R package to find and visualize monophyly issues. PeerJ Comput. Sci. 2, e56. 10.7717/peerj-cs.56.

Sakamoto, T., Ortega, J.M., 2020. TaxOnTree: a tool that generates trees annotated with taxonomic information. bioRxiv [preprint]. 10.1101/2020.12.24.424364.

Deng, Z., Botas, J., Cantalapiedra, C.P., Hernández-Plaza, A., Burguet-Castell, J., Huerta-Cepas, J., 2022. PhyloCloud: an online platform for making sense of phylogenomic data. Nucleic Acids Res. 50 (W1), W577–W582. 10.1093/nar/gkac324.

Sanderson, T., 2022. Taxonium, a web-based tool for exploring large phylogenetic trees. eLife 11, e82392. 10.7554/eLife.82392.

Letunic, I., Bork, P., 2021. Interactive Tree Of Life (iTOL) v5: an online tool for phylogenetic tree display and annotation. Nucleic Acids Res. 49 (W1), W293–W296. 10.1093/nar/gkab301.

Drummond, A.J., Rambaut, A., 2007. BEAST: Bayesian evolutionary analysis by sampling trees. BMC Evol. Biol. 7, 214. 10.1186/1471-2148-7-214.

Felsenstein, J., 1989. PHYLIP — Phylogeny Inference Package (Version 3.2) [software]. Cladistics 5, 164–166.

Bolyen, E., Rideout, J.R., Dillon, M.R., Bokulich, N.A., Abnet, C.C., Al-Ghalith, G.A., Alexander, H., Alm, E.J., Arumugam, M., Asnicar, F., Bai, Y., Bisanz, J.E., Bittinger, K., Brejnrod, A., Brislawn, C.J., Brown, C.T., Callahan, B.J., Caraballo-Rodríguez, A.M., Chase, J., Cope, E.K., Da Silva, R., Diener, C., Dorrestein, P.C., Douglas, G.M., Durall, D.M., Duvallet, C., Edwardson, C.F., Ernst, M., Estaki, M., Fouquier, J., Gauglitz, J.M., Gibbons, S.M., Gibson, D.L., Gonzalez, A., Gorlick, K., Guo, J., Hillmann, B., Holmes, S., Holste, H., Huttenhower, C., Huttley, G.A., Janssen, S., Jarmusch, A.K., Jiang, L., Kaehler, B.D., Kang, K.B., Keefe, C.R., Keim, P., Kelley, S.T., Knights, D., Koester, I., Kosciolek, T., Kreps, J., Langille, M.G.I., Lee, J., Ley, R., Liu, Y.-X., Loftfield, E., Lozupone, C., Maher, M., Marotz, C., Martin, B.D., McDonald, D., McIver, L.J., Melnik, A.V., Metcalf, J.L., Morgan, S.C., Morton, J.T., Naimey, A.T., Navas-Molina, J.A., Nothias, L.F., Orchanian, S.B., Pearson, T., Peoples, S.L., Petras, D., Preuss, M.L., Pruesse, E., Rasmussen, L.B., Rivers, A., Robeson, M.S., Rosenthal, P., Segata, N., Shaffer, M., Shiffer, A., Sinha, R., Song, S.J., Spear, J.R., Swafford, A.D., Thompson, L.R., Torres, P.J., Trinh, P., Tripathi, A., Turnbaugh, P.J., Ul-Hasan, S., van der Hooft, J.J.J., Vargas, F., Vázquez-Baeza, Y., Vogtmann, E., von Hippel, M., Walters, W., Wan, Y., Wang, M., Warren, J., Weber, K.C., Williamson, C.H.D., Willis, A.D., Xu, Z.Z., Zaneveld, J.R., Zhang, Y., Zhu, Q., Knight, R., Caporaso, J.G., 2019. Reproducible, interactive, scalable and extensible microbiome data science using QIIME 2. Nat. Biotechnol. 37, 852–857. 10.1038/s41587-019-0209-9.

MacIver, D.R., Hatfield-Dodds, Z., 2019. Hypothesis: a new approach to property-based testing. J. Open Source Softw. 4 (43), 1891. 10.21105/joss.01891.

Parks, D.H., Chuvochina, M., Waite, D.W., Rinke, C., Skarshewski, A., Chaumeil, P.-A., Hugenholtz, P., 2018. A standardized bacterial taxonomy based on genome phylogeny substantially revises the tree of life. Nat. Biotechnol. 36, 996–1004. 10.1038/nbt.4229.

Rinke, C., Chuvochina, M., Mussig, A.J., Chaumeil, P.-A., Davín, A.A., Waite, D.W., Whitman, W.B., Parks, D.H., Hugenholtz, P., 2021. A standardized archaeal taxonomy for the Genome Taxonomy Database. Nat. Microbiol. 6, 946–959. 10.1038/s41564-021-00918-8.

Parks, D.H., Chuvochina, M., Rinke, C., Mussig, A.J., Chaumeil, P.-A., Hugenholtz, P., 2022. GTDB: an ongoing census of bacterial and archaeal diversity through a phylogenetically consistent, rank normalized and complete genome-based taxonomy. Nucleic Acids Res. 50 (D1), D785–D794. 10.1093/nar/gkab776.

Parks, D.H., Chaumeil, P.-A., Mussig, A.J., Rinke, C., Chuvochina, M., Hugenholtz, P., 2026. GTDB release 10: a complete and systematic taxonomy for 715 230 bacterial and 17 245 archaeal genomes. Nucleic Acids Res. 54, gkaf1040. 10.1093/nar/gkaf1040.

Genome Taxonomy Database (GTDB), 2026. GTDB Release R232 [dataset]. https://gtdb.ecogenomic.org (accessed July 2026).

Kelly, K.L., 1965. Twenty-two colors of maximum contrast. Color Eng. 3 (6), 26–27.

Okabe, M., Ito, K., 2008. Color Universal Design (CUD) — how to make figures and presentations that are friendly to colorblind people. J*FLY. https://jfly.uni-koeln.de/color (accessed July 2026).

Zeng, Z., Wang, Y., 2026. FigTreeKit: A Python toolkit for programmatic FigTree styling, taxonomy-aware clade auditing, and rendering of phylogenetic trees (Version 1.1.1) [software]. Zenodo. 10.5281/zenodo.22106051.

Moody, E.R.R., Williams, T.A., Álvarez-Carretero, S., Szöllősi, G.J., Pisani, D., Lenton, T.M., Donoghue, P.C.J., 2025. The emergence of metabolisms through Earth history and implications for biospheric evolution. Philos. Trans. R. Soc. B 380, 20240097. 10.1098/rstb.2024.0097.

