## Supporting Information for "FigTreeKit: A Python toolkit for programmatic FigTree styling, taxonomy-aware clade auditing, and phylogenetic tree rendering"

### Supplementary Materials

#### Supplementary Section S1: Behavioral Specifications

##### S1.1 MRCA search

Precondition: the target taxa are strings expected among the tree tips. Postcondition (any-missing-fails contract, fix G1): _find_mrca_clade(tree, taxon_names) resolves the MRCA only if ALL requested taxa are present; if any target is absent the query returns None with a CompatibilityWarning listing the missing taxa, so a missing taxon can never produce a false monophyly verdict or an implicit collapse of the surviving subset. Partial-subset resolution is available solely through the explicit opt-in allow_missing=True, which is restricted to internal paths where targets may have been legitimately removed by a prior collapse; topology-gated collapse and monophyly queries never use it. collapse_clade() refuses requests with absent taxa unless allow_partial=True is passed explicitly, and check_monophyly() reports such requests as mrca_found=False. Tests: test/test_figtreekit.py::TestMRCAContractG1, test/test_taxonomy_correctness.py::TestRootingDependence.

##### S1.2 Iterative node-depth calculation

Precondition: a rooted Bio.Phylo tree. Postcondition: _calculate_node_height(node) returns the cumulative root-to-node branch length (depth); _get_min_tip_height(node) returns jebl's time-backward minimum tip height, which depends on the farthest tip of the subtree. The two semantics are separate, individually tested functions; neither is called "height" ambiguously in the API. Repeated queries are memoized per (tree, node) with a structural fingerprint guard (fix C6): cached entries are invalidated automatically after topology, rooting, or branch-length changes, and failure fallbacks (height 0.0 with CompatibilityWarning) are never cached; q uncached queries can require O(qn) time. O(n) time per query; O(n) stack worst case. Tests: test/test_conformance.py::TestNonUltrametricNodeHeight (pinned on a non-ultrametric counter-example tree).

##### S1.3 State-machine translate-block parsing

Precondition: a BEAST-style translate block. Postcondition: the parser is an explicit four-state machine (NORMAL, IN_SINGLE_QUOTE, IN_DOUBLE_QUOTE, ESCAPING); commas delimit entries only in NORMAL; doubled-quote escapes are resolved in ESCAPING; quoted names may contain commas and quotes of the other kind. Tests: test/test_conformance.py::TestTranslateRoundTrip (incl. escaped and comma-containing names).

##### S1.4 Position-aware multi-tree replacement

Precondition: a Nexus trees block with one or more tree declarations. Postcondition: tree boundaries are located by a character scanner tracking quote state and bracket-comment depth (arbitrary nesting), so semicolons inside quoted tree names or comments never terminate a tree; replacement affects only the selected tree_index span. The complete declaration grammar (tree <NAME> = <newick>; with quoted or unquoted names) and lexical precedence (quote state > bracket depth > semicolon) are documented in Code S4.5. Terminal-state validation (fix G2): in strict mode (default) the scanner raises ValueError for unmatched closing brackets, unterminated quotations or comments, and declarations lacking a terminating semicolon, and the serializer surfaces these as typed ExportError; a lenient mode preserves the historical salvage behavior. Tests: test/test_conformance.py::TestMultiTreeReplacementConformance, test/test_conformance.py::TestTreeDeclarationScannerValidation, test/test_rendering_mock.py::TestNestedBracketTreeDecl.

##### S1.5 Bracket-comment preservation

Precondition: any Newick/Nexus input with bracket comments. Postcondition (position support matrix): comments attached after tip names, after internal nodes, and after branch lengths are preserved through the Clade.comment round-trip and merged with injected attributes; root-level attributes ([&R]) are preserved on the root clade; if any comment is nevertheless lost, an explicit CompatibilityWarning is emitted — no silent loss. See Table S8. Tests: test/test_conformance.py::TestCommentPositionMatrix.

##### S1.6 Annotation injection consistency

Precondition: arbitrary interleaving of collapse and highlight calls. Postcondition (fix G3): highlight geometry and tip counts are computed from an immutable snapshot of the topology taken before any display collapse is applied; collapse annotations are then registered in deterministic inner-to-outer order. This sequence — snapshot first, collapse after — preserves pre-collapse descendant counts while producing the final collapsed display; the operation order and node-identity mapping are covered by explicit regression fixtures. Tests: test/test_figtreekit.py (hilight/collapse interleaving regressions).

##### S1.7 Serialization correctness

Precondition: any Python scalar used as a FigTree attribute value. Postcondition: booleans serialize as lowercase true/false; colors as lowercase unquoted hex; integer-valued floats without decimal point; strings double-quoted with backslash escaping. Tests: test/test_conformance.py::TestAnnotationFormatGolden, test/test_api_contracts.py.

##### S1.8 Monophyly detection

Precondition: a rooted tree and a taxonomy mapping; verdicts are qualified as "among mapped sampled tips", where "monophyletic" denotes recovery as an exclusive clade in the supplied rooted topology and does not establish biological support. The completeness audit is executed with the same mapping and parsing configuration before any verdict is interpreted. Postcondition: a group G is declared monophyletic iff the terminal set under MRCA(mapped members of G) equals that member set; unmapped tips nested inside the MRCA count as intruders and refuse the collapse; non-monophyletic groups are skipped with a warning. Tests: test/test_taxonomy_correctness.py (all six scenario classes).

**S1.9 Deep input validation**

Precondition: any input file offered to load/validate. Postcondition: bracket depth is non-negative everywhere and returns to zero; tip names are unique; branch lengths are non-negative in strict mode; FASTA/FASTQ alphabets are valid. Violations raise ValidationError. Tests: test/test_validators.py.

**S1.10 Tree-sequence cross-validation**

Postcondition: after cross-validation, tip set T and sequence ID set S satisfy $T \subseteq S$ and $S \subseteq T$ (mismatches reported explicitly). Tests: test/test_validators.py.

**S1.11 Anomalous-content screening (input hygiene)**

Postcondition: accepted content contains only safe printable code points; control characters and BiDi overrides (U+202A–202E, U+2066–2069) are rejected. This is input hygiene screening, not a security boundary. Tests: test/test_validators.py.

### Supplementary Section S2: Key Parameter Reference

**Table S1.** Key API parameters (units, defaults, allowed values).

| Parameter | Type | Default | Description |
|---|---|---|---|
| layout_type | LayoutType | RECTILINEAR | Tree layout: RECTILINEAR, POLAR, or RADIAL |
| branch_line_width | float (px) | 1.0 | Branch line width |
| font_size | int (pt) | 12 | Label font size |
| font_name | str | "Arial" | Font family name |
| font_style | int | 0 | 0=plain, 1=bold, 2=italic, 3=bold-italic |
| tree_index | int | 0 | Tree index to process in multi-tree files |
| strict | bool | False | Reject negative branch lengths / non-monophyletic collapses |
| max_taxa_warning_threshold | int | 10000 | Large-tree warning threshold |
| taxonomy_delimiter_mode | str | "reverse" | Taxonomy parsing: reverse / greedy / segment |
| taxonomy_source_priority | str | "table" | Priority: embedded vs. table taxonomy |
| taxonomy_table_sep | str | ";" | Format B rank separator |
| angular_range | float (deg) | 360.0 | Polar layout angular range |
| root_angle | float (deg) | 0.0 | Polar layout root angle |
| render_width | int (px) | 1200 | Render width |
| render_height | int (px) | 800 | Render height |
| collapse_type | str | "collapse" | "collapse" (triangle + label) or "cartoon" (triangle over tip range) |
| timeout | int (s) | 120 | JVM rendering wall-clock limit; raises RenderError on expiry |

**Table S2.** Command-line specific parameters (via --set KEY=VALUE).

| Key | Type | Description |
|---|---|---|
| tipLabels.isShown | bool | Show/hide tip labels |
| tipLabels.fontSize | int | Tip label font size |
| nodeLabels.isShown | bool | Show/hide node labels |
| scaleAxis.isShown | bool | Show/hide scale axis |
| scaleAxis.reverseAxis | bool | Reverse axis direction |
| polarLayout.angularRange | int (×1000) | Angular range (×1000) |
| polarLayout.rootAngle | int (×1000) | Root angle (×1000) |
| appearance.branchLineWidth | float | Branch line width |
| appearance.backgroundColour | color | Background color |

### Supplementary Section S3: Robustness Test Scenarios

**Table S3.** FigTreeKit robustness-test edge cases (28 scenarios).

| # | Edge case | Test result |
|---|---|---|
| 1 | Trifurcation (unrooted trifurcation) | Pass |
| 2 | Polytomy (10 children) | Pass |
| 3 | Tree with missing branch lengths | Pass |
| 4 | Taxon names with special characters inside quotes | Pass |
| 5 | Zero branch length | Pass |
| 6 | Negative branch length (strict mode) | Correctly rejected |
| 7 | BEAST translate block | Pass |
| 8 | Multi-tree Nexus file | Pass |
| 9 | Newick multi-tree file with --multi-tree | Correctly rejected |
| 10 | Legacy Java decimal colors | Pass |
| 11 | Floating-point precision (1,200-level deep tree) | Pass |
| 12 | Nexus with semicolons inside bracket comments | Pass |
| 13 | Quoted taxon names containing commas | Pass |
| 14 | Taxon names starting with numbers (e.g., 9606) | Pass |
| 15 | Taxonomy embedded format A | Pass |
| 16 | Taxonomy table format B | Pass |
| 17 | Monophyletic collapsing | Pass |
| 18 | Nested collapsing (inner (A,B) + outer (A,B,C)) | Pass |
| 19 | Collapse label assigned to representative node | Pass |
| 20 | Non-monophyletic collapsing (strict mode) | Correctly rejected |
| 21 | PNG image rendering | Pass |
| 22 | PDF vector rendering | Pass |
| 23 | SVG vector rendering | Pass |
| 24 | Anomalous control characters | Correctly rejected |
| 25 | FASTA @ lines not misclassified as FASTQ | Pass |
| 26 | Tree-sequence cross-validation | Pass |
| 27 | set_clade_color + highlight_clade conflict | CompatibilityWarning issued |
| 28 | _hilight_marks instance isolation | Pass |

### Supplementary Section S4: Core Algorithm Implementation Code

##### Code S4.1 MRCA search algorithm

```
def _find_mrca_clade(self, tree, taxon_names, allow_missing=False):
    # MRCA contract (G1): ALL requested taxa must be present before
    # resolution; partial resolution is an explicit opt-in used only by
    # internal paths that legitimately handle collapse-removed targets.
    try:
        names = list(taxon_names)
        tips = {t.name for t in tree.get_terminals()}
        missing = [n for n in names if n not in tips]
        if missing and not allow_missing:
            warnings.warn(f"MRCA search failed for taxa {taxon_names}: "
                          f"{len(missing)} target(s) absent; refusing "
                          f"partial resolution.", CompatibilityWarning)
            return None
        if missing:                      # explicit partial mode
            names = [n for n in names if n in tips]
        if not names:
            warnings.warn(f"MRCA search failed for taxa {taxon_names}: "
                          f"no matching taxa found in tree")
            return None
        if len(names) == 1:
            for t in tree.get_terminals():
                if t.name == names[0]:
                    return t
            return None
        return tree.common_ancestor(names)
    except (ValueError, AttributeError, KeyError) as e:
        warnings.warn(f"MRCA search failed for taxa {taxon_names}: {e}")
        return None
```

##### Code S4.2 MRCA resolution from node references (nested collapsing)

```
@staticmethod
def _find_mrca_of_nodes(tree, nodes):
    if not nodes:
        return None
    if len(nodes) == 1:
        return nodes[0]
    mrca = nodes[0]
    for node in nodes[1:]:
        mrca = tree.common_ancestor(mrca, node)
    return mrca
```

##### Code S4.3 Iterative node-depth calculation (root-to-node depth)

```
def _calculate_node_height(self, tree, node) -> float:
    # Memoized per (tree, node) with a structural guard (C6): entries are
    # dropped when the tree's fingerprint (id, terminal count, total branch
    # length) changes; failure fallbacks are deliberately not cached.
    cache = self.__dict__.setdefault('_node_height_cache', {})
    key = (id(tree), id(node))
    cached = cache.get(key)
    if cached is not None:
        result, guard = cached
        if guard == self._tree_cache_guard(tree):
            return result
        del cache[key]                   # stale entry: tree mutated
    stack, found = [(tree.root, 0.0)], None
    while stack:                         # iterative DFS, no recursion
        current, height = stack.pop()
        if current is node:
            found = round(height, 10)
            break
        for child in getattr(current, 'clades', []):
            stack.append((child, height + (child.branch_length or 0.0)))
    if found is None:
        warnings.warn("Could not find path from root to node; "
                      "height defaults to 0.0", CompatibilityWarning)
        return 0.0                       # not cached
    cache[key] = (found, self._tree_cache_guard(tree))
    return found
```

**Code S4.4 Four-state-machine translate-block parsing**

```
state = TRANSLATE_NORMAL   # NORMAL / IN_SINGLE_QUOTE / IN_DOUBLE_QUOTE / ESCAPING
open_quote = ''
for char in content:
    if state == TRANSLATE_ESCAPING:
        if char == open_quote:
            current += char        # doubled quote: one escaped literal
            state = (TRANSLATE_IN_SINGLE_QUOTE if open_quote == "'"
                     else TRANSLATE_IN_DOUBLE_QUOTE)
            continue
        state = TRANSLATE_NORMAL   # closing quote; reprocess char below
    if state == TRANSLATE_NORMAL:
        if char == "'":
            state, open_quote = TRANSLATE_IN_SINGLE_QUOTE, char
            current += char
        elif char == '"':
            state, open_quote = TRANSLATE_IN_DOUBLE_QUOTE, char
            current += char
        elif char == ',':
            entries.append(current.strip()); current = ''
        else:
            current += char
    else:  # IN_SINGLE_QUOTE or IN_DOUBLE_QUOTE
        matching = "'" if state == TRANSLATE_IN_SINGLE_QUOTE else '"'
        if char == matching:
            open_quote = matching
            state = TRANSLATE_ESCAPING
            current += char
        else:
            current += char
# EOF handling (C5): validate the terminal state before appending the
# final entry; doubled-quote decoding keeps exactly one literal quote.
if state in (TRANSLATE_IN_SINGLE_QUOTE, TRANSLATE_IN_DOUBLE_QUOTE):
    raise ValidationError("Unterminated quote in translate block")
last = current.strip()
if last:
    if re.match(r'^\d+$', last.split(None, 1)[0]) is None:
        raise ValidationError(f"Malformed final translate entry: {last!r}")
    entries.append(last)
```

**Code S4.5 Position-aware multi-tree replacement (character scanner)**

```
def find_tree_declaration_spans(trees_content, strict=True):
    # Grammar: tree <NAME> = <newick>;  NAME = quoted ('' / "" escaping)
    # or unquoted non-whitespace token. Lexical precedence:
    # quote state > bracket-comment depth > tree-delimiting semicolon.
    spans, n, search_from = [], len(trees_content), 0
    while True:
        decl = _TREE_NAME_PATTERN.search(trees_content, search_from)
        if decl is None:
            break
        j, in_quote, bracket_depth = decl.end(), None, 0
        while j < n:
            char = trees_content[j]
            if in_quote is not None:
                if char == in_quote:
                    if j + 1 < n and trees_content[j + 1] == in_quote:
                        j += 2; continue       # doubled-quote escape
                    in_quote = None
            elif char in ("'", '"'):
                in_quote = char
            elif char == '[':
                bracket_depth += 1
            elif char == ']':
                if bracket_depth == 0 and strict:
                    raise ValueError("Unmatched closing bracket ']'")
                bracket_depth = max(0, bracket_depth - 1)
            elif char == ';' and bracket_depth == 0:
                break
            j += 1
        if j >= n:                     # terminal-state validation (G2)
            if not strict:             # legacy salvage mode
                spans.append((decl.start(), n)); break
            if in_quote is not None:
                raise ValueError("Unterminated quotation")
            if bracket_depth > 0:
```

```
            raise ValueError("Unterminated bracket comment")
        raise ValueError("Missing terminating semicolon")
    spans.append((decl.start(), j + 1))
    search_from = j + 1
return spans
```

##### Code S4.6 Bracket-comment preservation mechanism

```
def _resolve_annotations_copy(self):
    # Trees are parsed WITHOUT stripping bracket comments: Biopython >= 1.80
    # round-trips them via Clade.comment, so BEAST metadata survives at every
    # node-level position; injected attributes merge into existing comments.
    bracket_comments = self._extract_bracket_comments(self._tree_content)
    tree = self._parse_tree_with_biopython(self._tree_content)
    resolved_annotations = self._build_resolved_annotations()
    self._apply_annotations_to_tree(tree, resolved_annotations)
    serialized = self._serialize_tree_to_newick(tree)
    if bracket_comments:
        # Re-insertion is guarded: comments already present (possibly merged)
        # are skipped; genuinely lost positions emit CompatibilityWarning.
        serialized = self._reinsert_bracket_comments(serialized, bracket_comments)
    return serialized
```

##### Code S4.7 FigTree command-line rendering (error-checked)

```
result = subprocess.run(cmd, capture_output=True, text=True, timeout=timeout)
success = (result.returncode == 0
           and os.path.isfile(output_file)
           and os.path.getsize(output_file) > 0)
if success:
    return True
if result.returncode != 0:
    raise RenderError(f"FigTree exited with code {result.returncode}\n{detail}")
if not os.path.isfile(output_file):
    raise RenderError(f"FigTree rendering produced no output file: {output_file}")
if os.path.getsize(output_file) == 0:
    raise RenderError("FigTree rendering produced an empty output file")
# subprocess.TimeoutExpired is likewise converted to RenderError.
```

### Supplementary Section S5: FigTreeStyler Class Architecture

**Table S4.** FigTreeStyler class architecture.

| Layer | Responsibility | Example methods | Module |
|---|---|---|---|
| Loading/parsing | Read Newick/Nexus | load_tree(), load_content() | _parser.py |
| Styling | Layout, font, color | set_layout(), set_appearance() | styler.py |
| Annotation | Inject !hilight, !color, !font | highlight_clade(), set_clade_color_all() | styler.py + _serializer.py |
| Taxonomy | Parse, detect monophyly, collapse | analyze_taxonomy(), collapse_by_group() | taxonomy.py |
| Serialization | Write Nexus | export() | _serializer.py |
| Validation | Structural checks | validate() | validators.py |
| Rendering | FigTree CLI | render() | _renderer.py |

### Supplementary Section S6: Complexity Analysis

**Table S5.** Core algorithm complexity.

| Algorithm | Time | Space | Implementation |
|---|---|---|---|
| MRCA search (k targets) | O(k·n) | O(n) auxiliary (terminal-name set) + O(h) call stack | Bio.Phylo recursive (any-missing-fails contract, G1) |
| Node-depth calculation | O(n) | O(n) worst case; O(h) balanced | Iterative DFS (memoized with structural guard, C6) |
| Translate-block parsing | O(m) | O(k) | Four-state machine |
| Multi-tree replacement | O(L) | O(L) | Character scanner (arbitrary nesting) |
| Annotation injection (single) | O(n) | O(h) call stack | Bio.Phylo recursive |
| Taxonomy extraction | O(n) | O(n) | Regex/string |
| Monophyly detection (per group) | O(n) | O(n) | Bio.Phylo recursive |
| Nested-collapse sorting (k groups) | O(k·n + k log k) | O(k) | Sort + node-ref MRCA |
| Full export (a annotations) | O(a·n) | O(n) | Mixed — super-linear when a ∝ n |
| Image rendering | Empirical: JVM-startup constant + size/label/annotation-dependent term | Empirical (not bounded by O(r) alone, G4) | FigTree CLI (subprocess) |

*n = nodes, h = tree height, m = translate-block length, k = taxa/group count, g = nested-group count, L = file length, a = annotations. As implemented (fix G4): MRCA resolution allocates a terminal-name set and therefore requires O(n) auxiliary space in addition to the O(h) call-stack cost of the recursive Bio.Phylo traversal; recursive annotation traversals require O(h) call-stack space rather than O(1); per-group topology assessment inherits the target-dependent MRCA cost; sorting g nested groups adds O(g log g); and q uncached depth queries can require O(qn). Rendering cost depends on tree size, labels, annotations, vector complexity, and raster dimensions, and is characterized empirically (JVM startup ~0.8–1.2 s constant plus a size-dependent term) rather than by a single O(r) bound. The annotation sweep in results_annotations.csv confirms the O(a·n) term empirically.*

### Supplementary Section S7: Functional Comparison Evaluation Criteria

**Table S6.** Functional comparison evaluation criteria.

| Feature | Full support | Partial | Not supported |
|---|---|---|---|
| Newick/Nexus I/O | Read and write | Read-only or write-only | Neither |
| FigTree annotation | Generate the three supported annotation types (!hilight, !color, !font) | Some types only | Cannot generate |
| BEAST translate block | Correct handling + metadata preservation | Parse but lose metadata | Cannot parse |
| Programmatic styling | Full API + FigTree-format output | API exists, non-FigTree output | No API |
| Taxonomy analysis | Parsing + monophyly + collapse | Basic parsing only | No taxonomy |
| CLI rendering | Direct command-line export | Requires extra config | No CLI |
| Validation | Deep validation + hygiene screening | Basic checks only | No validation |

*Tool versions evaluated (latest stable releases available as of the July 2026 search date): Bio.Phylo 1.87, DendroPy 4.6.0, ETE3 3.1.3, ggtree 3.6.0, TreeViewer 2.2.0, FigTreeKit 1.1.1; phylotreelib, collapseGTDB, and figtree-recolor were evaluated from their public PyPI/GitHub revisions (Section S8, Figure S7). Each rating reflects the documented capabilities of the evaluated version; cases verified only from documentation rather than executed fixtures are marked explicitly, and untested cases are reported as unknown rather than inferred.*

### Supplementary Section S8: Tool-by-Tool Differentiation

**Table S7.** Tool-by-tool differentiation from adjacent software.

| Tool | Reference | What it does | Why it does not fill FigTreeKit's niche |
|---|---|---|---|
| MonoPhy | Schwery & O'Meara (2016) | R package to find and visualize monophyly issues | Monophyly assessment only; no FigTree styling annotations or programmatic layout/color/highlight control |
| TaxOnTree | Sakamoto & Ortega (2020) | Decorates internal tree nodes with taxonomic information for inspection in FigTree | Taxonomy labelling, not styling annotation; no programmatic access to !hilight/!color/!font |
| PhyloCloud | Deng et al. (2022) | Web platform for exploring phylogenomic data at scale | Online platform, not a scriptable Python library; no FigTree-compatible Nexus export |
| Taxonium | Sanderson (2022) | Web-based exploration of very large trees (>100,000 taxa) | Interactive viewer for a different scale regime; complementary (recommended in §4.1) |
| iTOL | Letunic & Bork (2021) | The most widely used online platform for tree display and annotation | iTOL-specific annotation files within its web ecosystem; cannot generate FigTree Nexus annotations |
| phylotreelib | https://pypi.org/project/phylotreelib/ | Python tree-format library; writes NEXUS with optional FigTree color attributes | Tip-color subset only; no layout, highlight, font, collapse, translate handling, or taxonomy-aware workflow |
| collapseGTDB | https://github.com/genomewalker/collapse-gtdb-tree | Collapses GTDB reference trees at a chosen taxonomic rank | Rank collapse without monophyly validation reporting, no FigTree styling annotations, no rendering integration |
| figtree-recolor | https://github.com/acorg/figtree-recolor | Script that rewrites tip colors in FigTree-saved Nexus files | Single annotation type on tips; self-described hack without tests; no batch/validation/taxonomy features |

*All tools are excellent within their own paradigms; this table concerns only the specific niche of programmatic generation of FigTree-compatible styling annotations combined with taxonomy-aware, monophyly-validated collapse. Exact FigTree 1.4.4 serialization is reported once as a specific interoperability target rather than as a general quality axis.*

### Supplementary Section S9: Correctness and Compatibility Validation

This section reports the golden conformance corpus, the comment-position support matrix, deterministic scenario-based verification with simulated ground truth, and real-data audits referenced by main-text Section 3.1/3.4.

**Table S8.** Bracket-comment position support matrix.

| Attachment position | Behavior | Guarantee |
|---|---|---|
| After tip taxon name | Preserved via Clade.comment round-trip | Observed in tested fixtures |
| After internal node ')' | Preserved; merges with injected attributes | Observed in tested fixtures |
| After branch length ':x' | Preserved; merges with injected attributes | Observed in tested fixtures |
| Root/block level ([&R]) | Root/block level ([&R], tree-level rooting directive) | Tested separately from node comments |
| Any position where loss still occurs | Explicit CompatibilityWarning | Explicit CompatibilityWarning; case listed in the machine-readable audit |

**Table S9**. Deterministic scenario-based verification of taxonomy-aware collapse on simulated trees with known ground truth (test_taxonomy_correctness.py). These scenarios verify expected behavior for selected topological and mapping edge cases; they constitute scenario-based verification rather than an accuracy benchmark and do not estimate false-positive or false-negative rates.

| Scenario | Expected | Observed |
|---|---|---|
| Monophyletic group | Detected; collapse registered | Detected; [&!collapse] emitted |
| Paraphyletic group | Refused; intruder taxa reported | Refused; intruder listed; 0 collapses |
| Polyphyletic group | Refused | Refused; 0 collapses |
| Clade inside polytomy | Detected | Detected |
| Same tips, correct vs. wrong rooting | Verdict flips with root | Monophyletic ↔ non-monophyletic |
| Unmapped tip nested inside MRCA | Refused (conservative) | Refused; unmapped tip reported as intruder |
| Unmapped tip outside MRCA | Monophyletic among mapped tips; audited | Verdict issued; unmapped list + completeness audit exposed |

**Table S10.** GTDB R232 ar53 workflow audit (examples/05_gtdb_workflow.py, same frozen run). Group accounting distinguishes assessed groups, multi-tip groups producing non-trivial collapses (142), and singleton orders that are trivially exclusive but leave the display unchanged (37); performance values correspond to the 2026-08-26 frozen run.

| Metric | Value |
|---|---|
| Mapped tips / total tips | 10,122 / 10,122 |
| Mapping coverage (completeness audit) | 100% |
| Phylum-level groups (monophyletic / non-monophyletic) | 25 / 0 |
| Order-level groups assessed (recovered as exclusive / skipped) | 179 (179 / 0) |
| Non-trivial collapses (multi-tip orders) / singleton no-ops | 142 / 37 (singleton collapses leave the display unchanged) |
| Parse / export time (2026-08-26 frozen run) | 0.06 s / 0.16 s |

**Table S11.** LACA BEAST real-data audit (aggregate checks). The table reports aggregate counts (input hash, version, and failure counts are recorded in the workflow output); the in→out equality of annotation counts demonstrates no loss at aggregate level, while clade-level key/value/association comparisons for the 699 retained annotations are provided by the machine-readable audit produced by examples/06_beast_laca_workflow.py.

| Metric | Value |
|---|---|
| Tips | 700 |
| [&95%={…}] annotations in → out (aggregate) | 699 → 699 (equal aggregate counts; clade-level key/value/association comparisons in the machine-readable audit) |
| Phylum-level groups (monophyletic / non-monophyletic) | 119 / 6 |
| Unmapped tips | 0 |
| Headless render (polar layout, PDF) | OK (patched JAR) |

#### Supplementary Figures

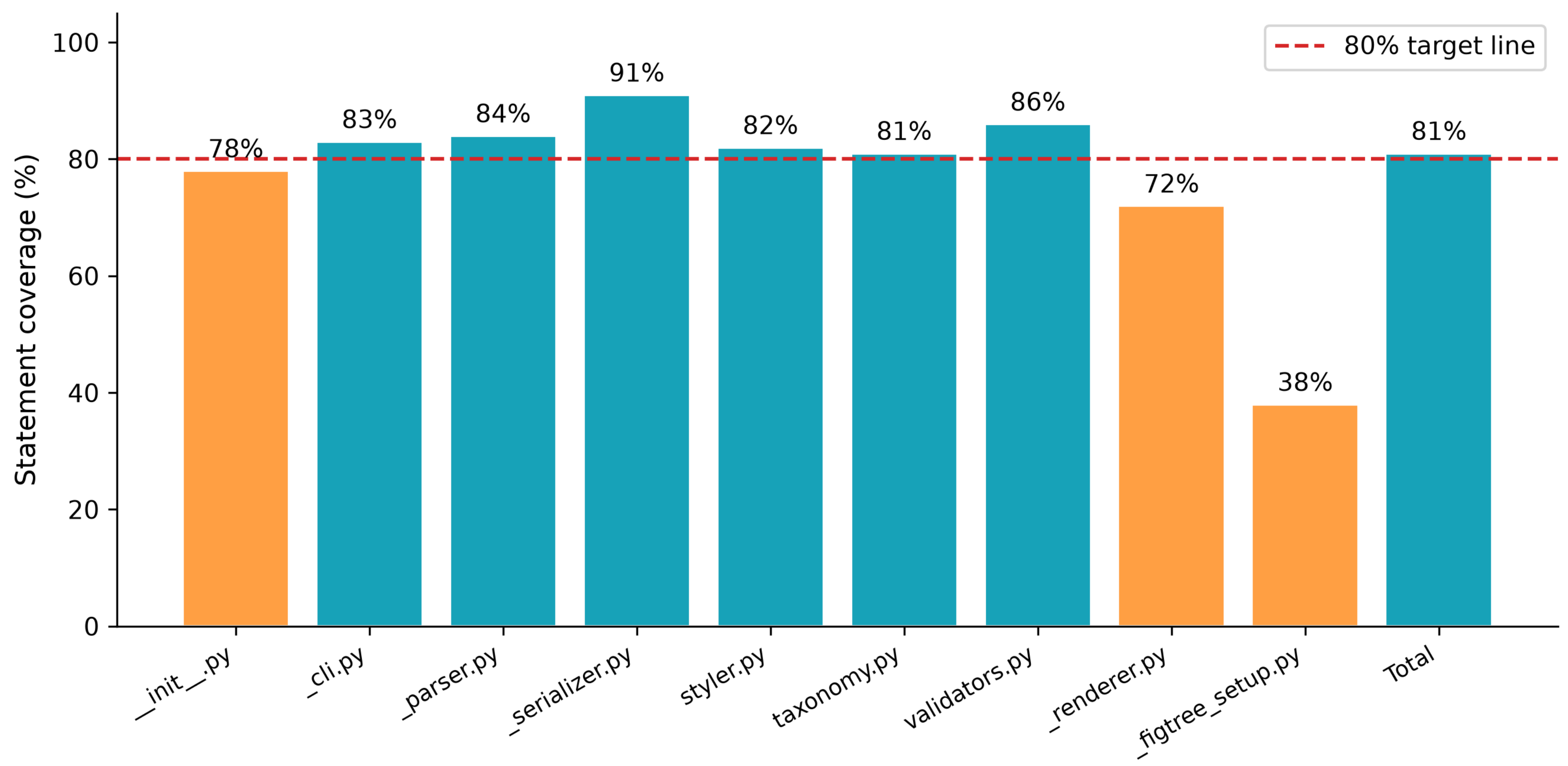


**Figure S1. Per-module statement coverage of the FigTreeKit test suite (796 passing tests).** Orange bars fall below the 80% target line. Coverage is reported as a descriptive engineering metric, not as evidence of correctness.

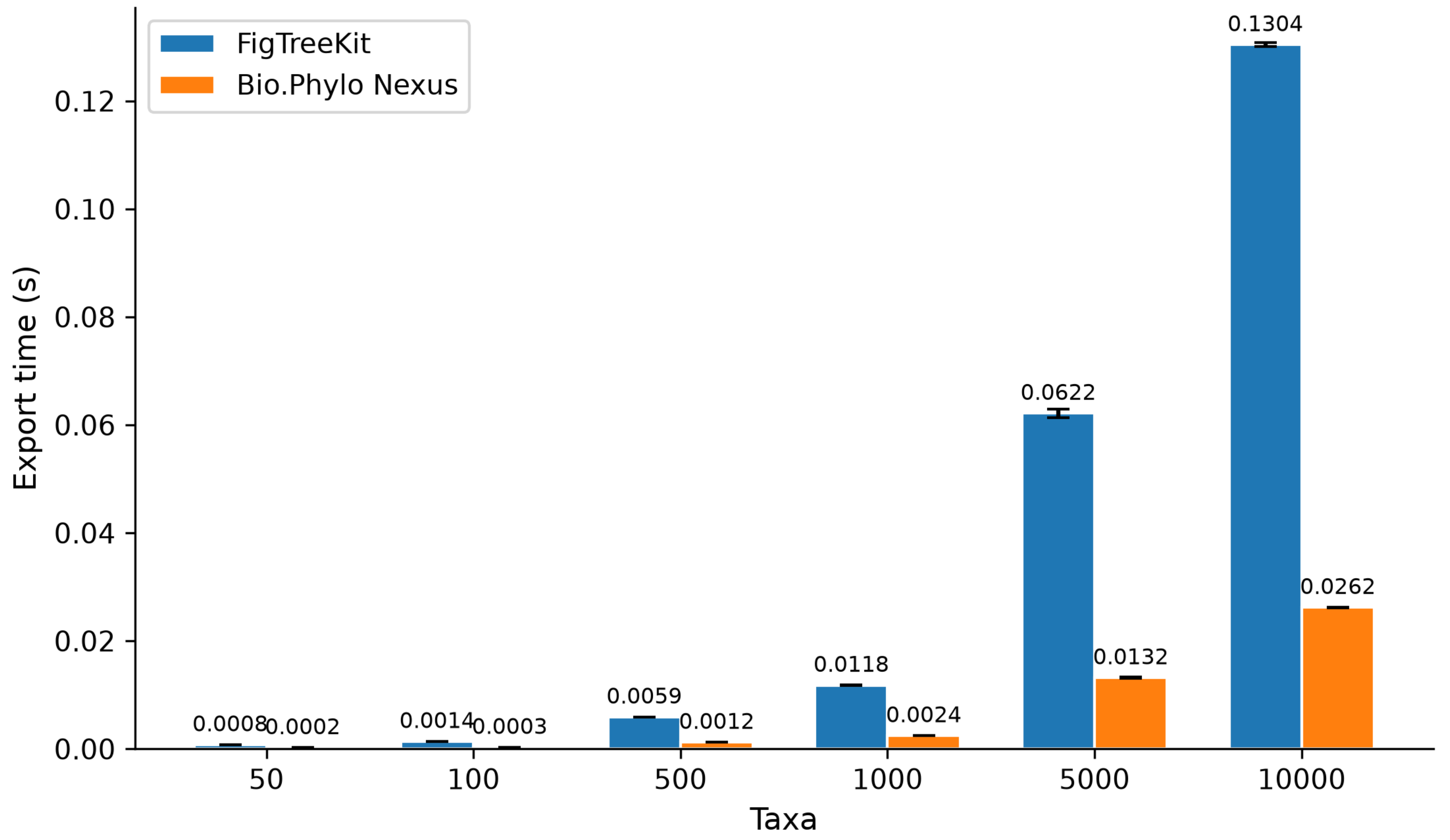


**Figure S2. Serialization-overhead microbenchmark: FigTreeKit versus Bio.Phylo NEXUS export (export-only, no annotation injection).** Paired measurements over 10 independently generated balanced trees per size (per-tree medians of 10 technical repeats); bars show tree-level mean ± SEM.

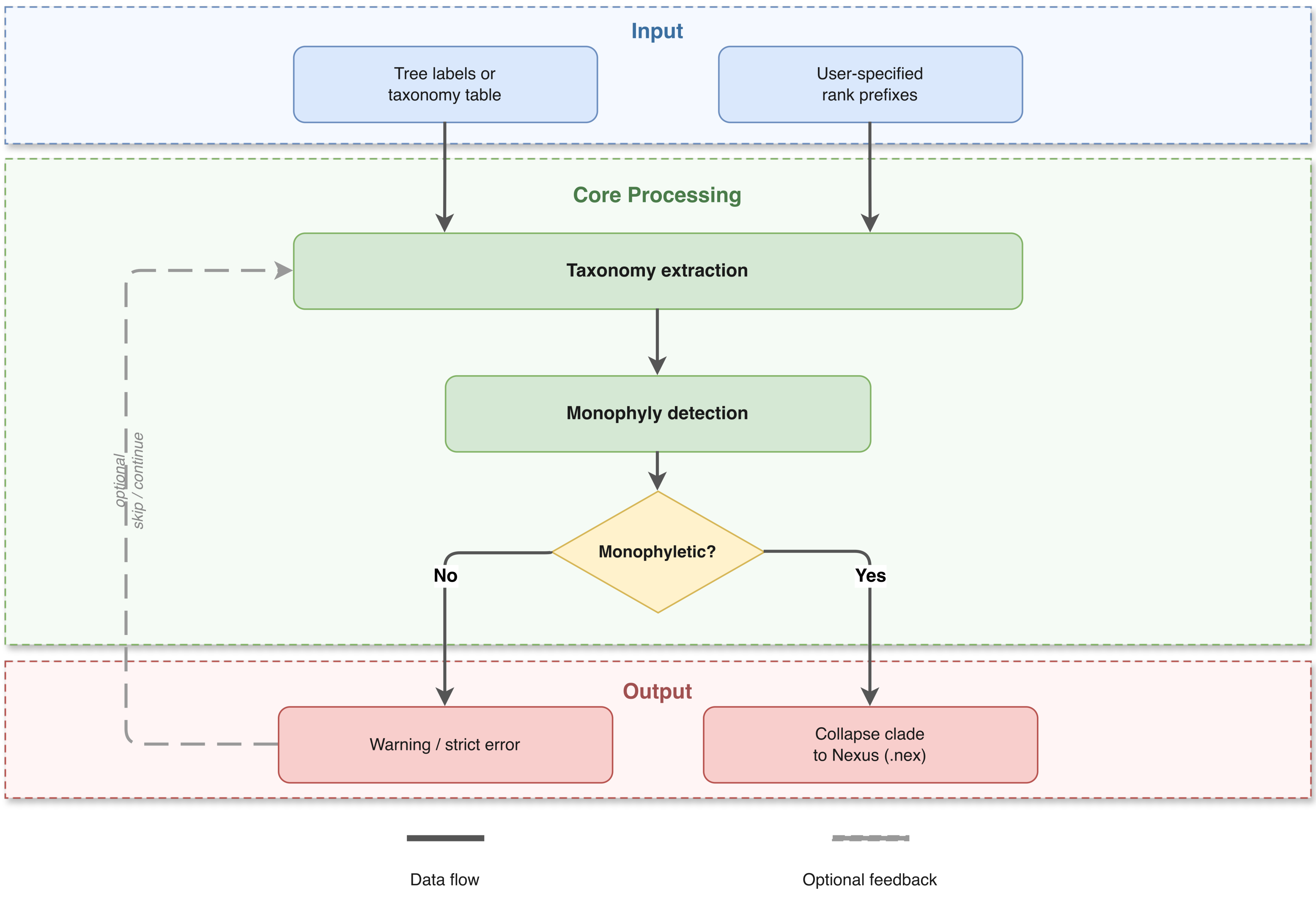


**Figure S3.** Taxonomy-aware analysis workflow: labels/mapping → TaxonomyMapper → MonophylyAnalyzer → validated color/collapse or skip-with-warning. Completeness auditing precedes every verdict.

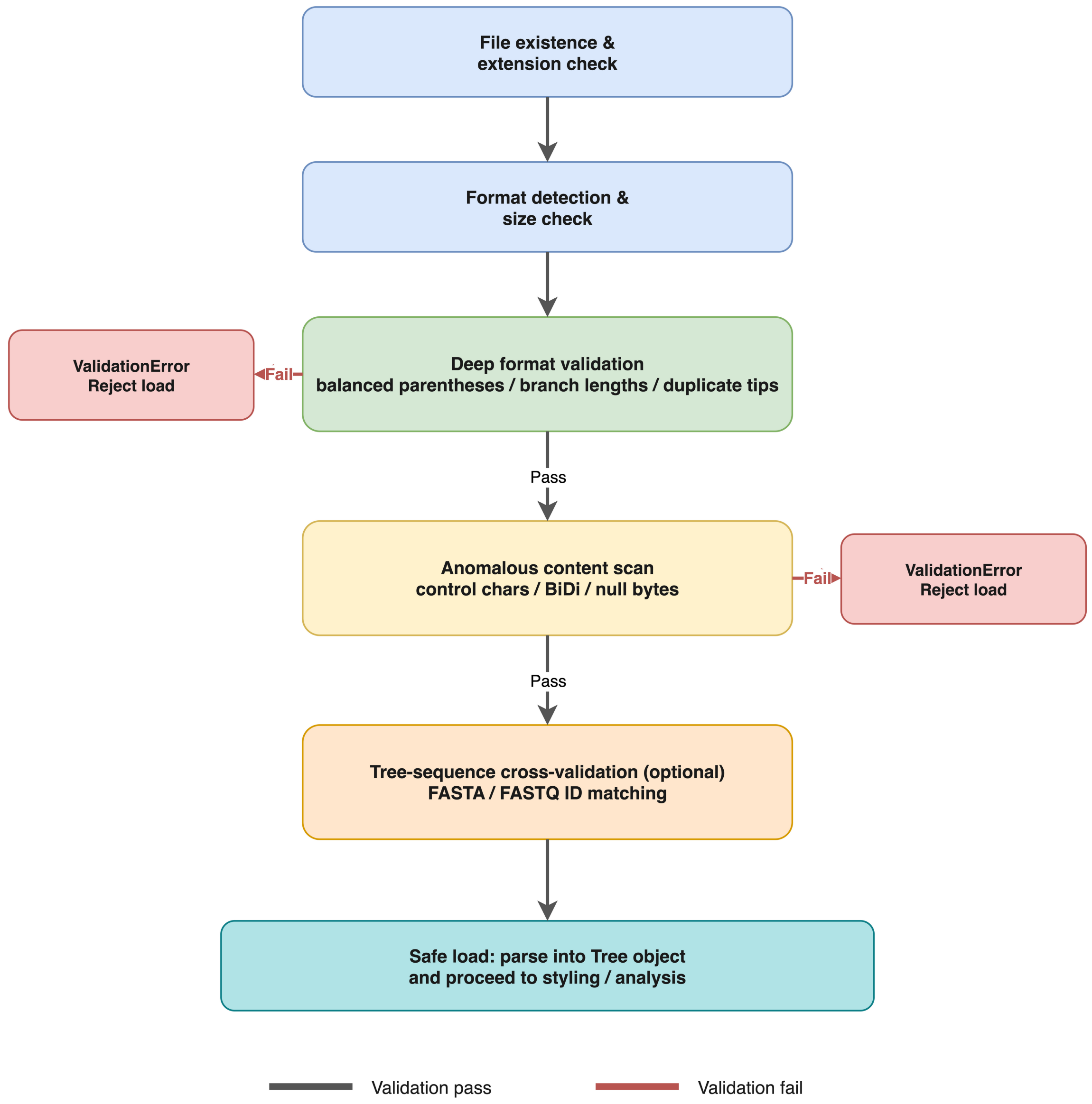


**Figure S4.** Input validation and hygiene screening workflow (format detection → structural validation → anomalous-character screening → accept/ValidationError + CompatibilityWarning).

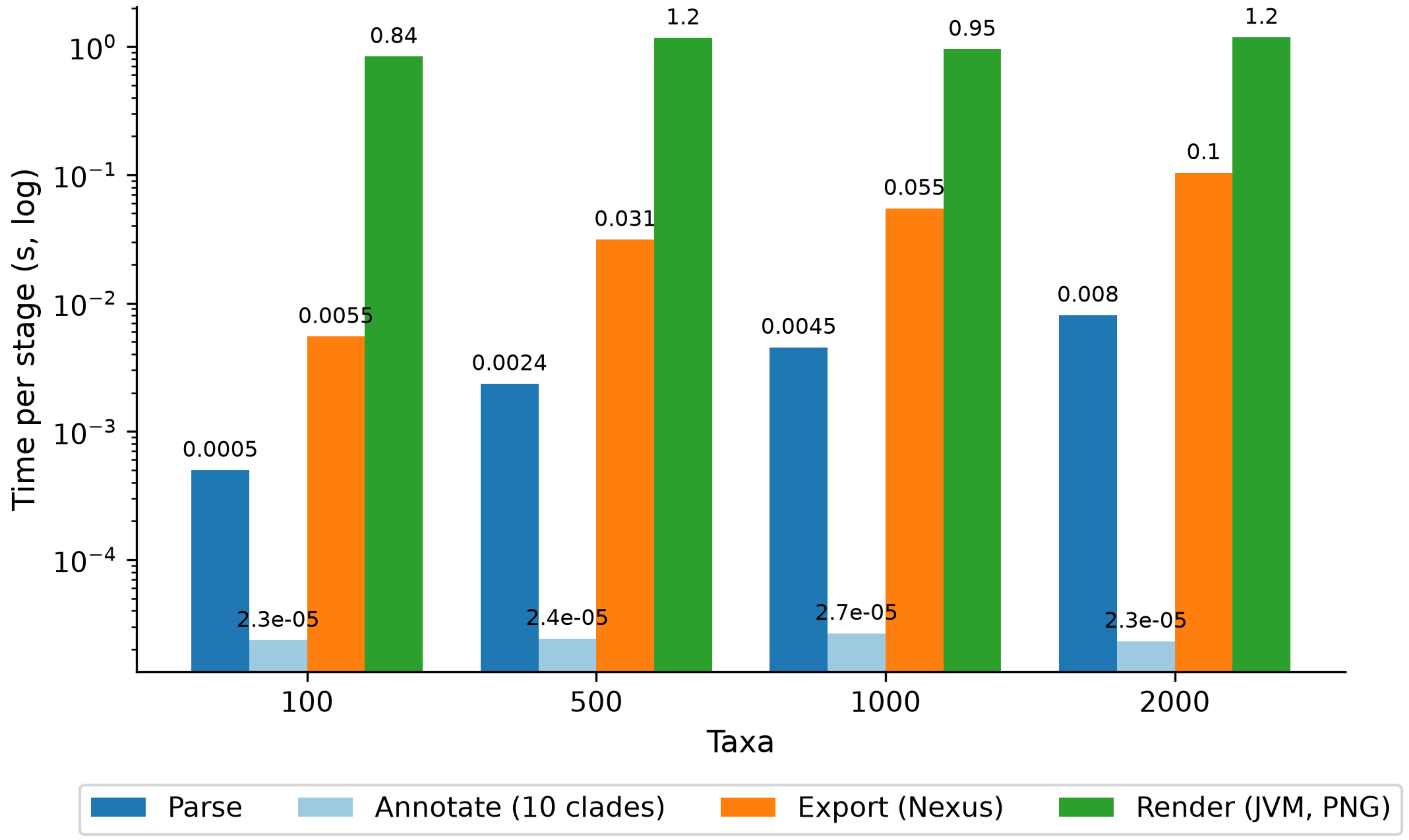


**Figure S5.** FigTreeKit pipeline stage breakdown (parse / annotate / export / render) for 100–2,000-taxon trees, median of 3 runs. JVM startup dominates rendering (~1.1–1.3 s constant), which is why per-tree render speed is not compared against native renderers in this work.

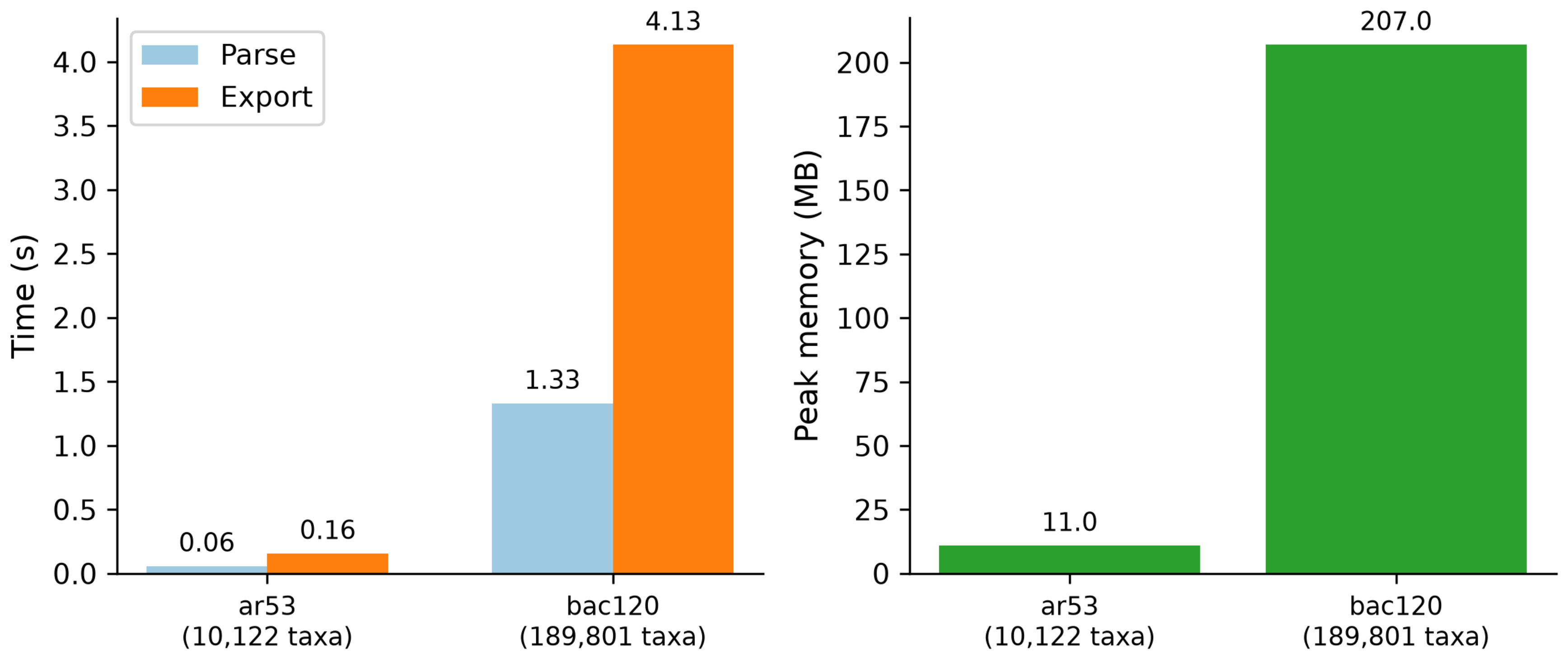


**Figure S6. GTDB R232 large-data scalability demonstration:** (A) parse/export times; (B) peak memory. ar53 (10,122 taxa): 0.06 s / 0.16 s, 11.0 MB; bac120 (189,801 taxa): 1.33 s / 4.14 s, 207.0 MB. Machine-stamped raw values: benchmarks/gtdb_results.json.

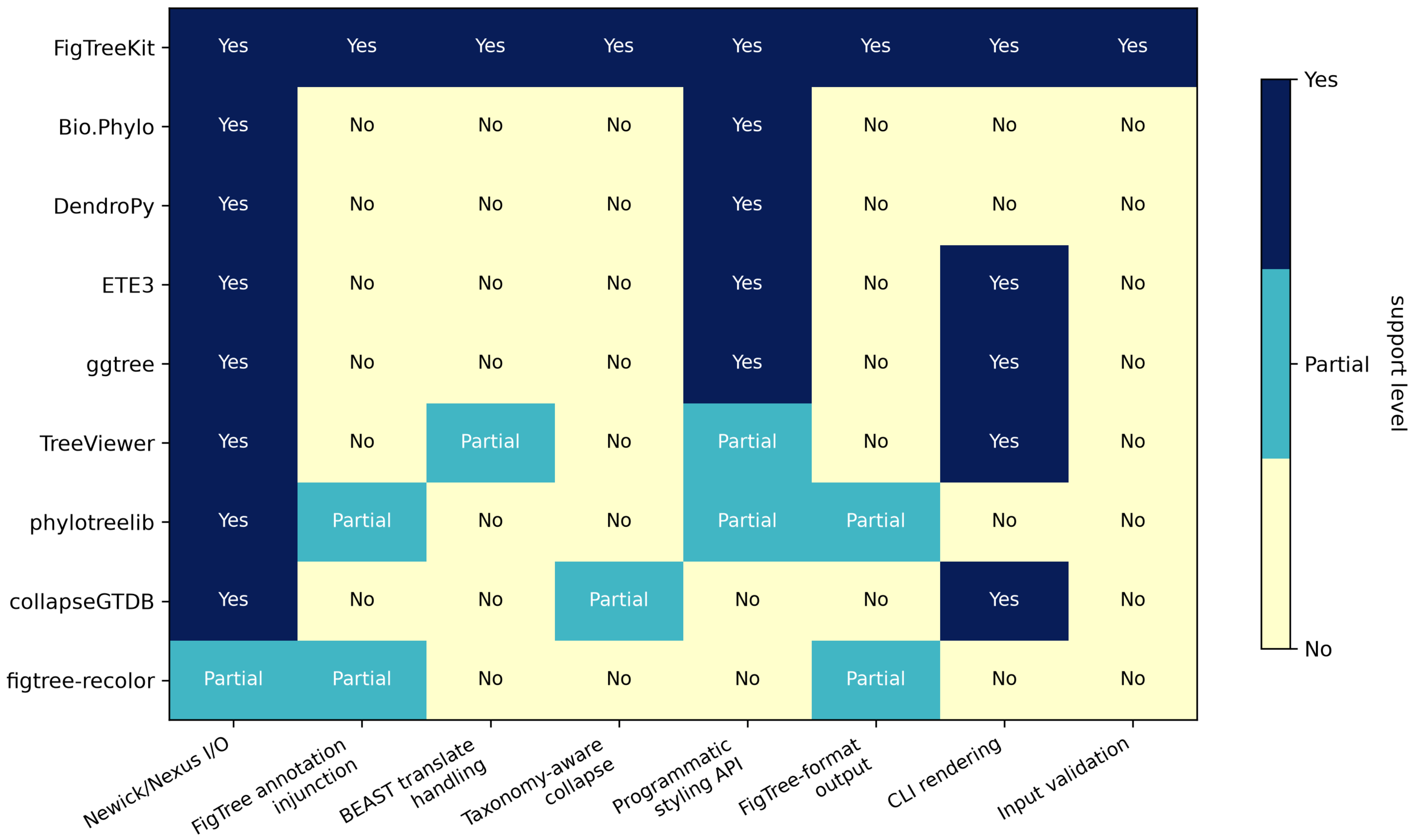


**Figure S7.** Feature matrix heat map — extended view of Table 1 including phylotreelib, collapseGTDB, and figtree-recolor. Scoring criteria: Table S6; tool-by-tool rationale: Table S7.
